# RAF inhibitors activate the integrated stress response by direct activation of GCN2

**DOI:** 10.1101/2024.08.15.607884

**Authors:** Rebecca Gilley, Andrew M. Kidger, Graham Neill, Paul Severson, Dominic P. Byrne, Niall S. Kenneth, Gideon Bollag, Chao Zhang, Taiana Maia de Oliveira, Patrick A. Eyers, Richard Bayliss, Glenn R. Masson, Simon J. Cook

## Abstract

Paradoxical RAF activation by chemical RAF inhibitors (RAFi) is a well-understood ‘on-target’ biological and clinical response. In this study, we show that a range of RAFi drive ERK1/2-independent activation of the Unfolded Protein Response (UPR), including expression of ATF4 and CHOP, that required the translation initiation factor eIF2α. RAFi-induced ATF4 and CHOP expression was not reversed by inhibition of PERK, a known upstream activator of the eIF2α-dependent Integrated Stress Response (ISR). Rather, we found that RAFi exposure activated GCN2, an alternate eIF2α kinase, leading to eIF2α-dependent (and ERK1/2-independent) ATF4 and CHOP expression. The GCN2 kinase inhibitor A-92, GCN2 RNAi, GCN2 knock-out or ISRIB (an eIF2α antagonist) all reversed RAFi-induced expression of ATF4 and CHOP indicating that RAFi require GCN2 to activate the ISR. RAFi also activated full-length recombinant GCN2 *in vitro* and in cells, generating a characteristic ‘bell-shaped’ concentration-response curve, reminiscent of RAFi-driven paradoxical activation of WT RAF dimers. Activation of the ISR by RAFi was abolished by GCN2 kinase dead mutations and M802A or M802G gatekeeper mutations, suggesting that RAFi bind directly to the GCN2 kinase domain; this was supported by mechanistic structural models of RAFi interaction with GCN2. Since the ISR is a critical pathway for determining cell survival or death, our observations may be relevant to the clinical use of RAFi, where paradoxical GCN2 activation may be a previously unappreciated off-target effect that may modulate tumour cell responses.

## Introduction

The RAS-regulated RAF-MEK1/2-ERK1/2 signalling pathway is activated in a variety of cancers due to mutations in RAS (especially KRAS), BRAF and more rarely CRAF, MEK1 or MEK2^1–4^. The most common BRAF mutations, BRAF^V600E/K^, are found in melanoma, hairy cell leukemia, thyroid cancer and colorectal cancer and signal as constitutively active monomers to drive MEK1/2-ERK1/2 activation and disease progression^5,6^. BRAF^V600E/K^ mutants are effectively inhibited by the clinically approved RAF inhibitors (RAFi) Vemurafenib, Dabrafenib and Encorafenib^3,7–9^, which are now used in combination with MEK1/2 inhibitors to treat BRAF^V600E/K^ mutant metastatic melanoma^10^ and colorectal cancer^11^.

In contrast, a variety of RAFi have consistently failed to progress through the clinic for tumours with wild type (WT) RAF, including those driven by RAS mutations, because they cause paradoxical activation of RAF and ERK1/2 signalling^12,13^. This is because WT RAF proteins signal as RAS-dependent homo- or heterodimers in which one protomer can transactivate the other. At sub-saturating doses RAFi promote dimerization^14–16^ and whilst binding of RAFi to one protomer inhibits it, this drives allosteric transactivation of the inhibitor-free, ATP-bound dimer partner, resulting in drug-induced activation of MEK1/2-ERK1/2 in cells. Furthermore, RAFi binding to one protomer of the dimer reduces the binding affinity and drug occupancy time of the second protomer^17^. This negative cooperativity means that only at high doses of RAFi are both dimer partners inhibited and ERK1/2 activation blocked; this accounts for RAFi eliciting a characteristic broad bell-shaped concentration-response curve for paradoxical ERK1/2 activation in cells containing WT RAF. This paradoxical RAF-MEK1/2-ERK1/2 activation limits efficacy in BRAF wild type tumours but also drives adventitious growth of low-grade tumours in tissue that lacks BRAF^V600E/K^ ^18^. RAFi-induced activation of MEK1/2-ERK1/2 can be mitigated by combination with a MEKi such as Trametinib; this combination enhances ERK1/2 inhibition in BRAF^V600E/K^ tumour cells but antagonises paradoxical ERK1/2 activation in tissue that lacks BRAF mutations, thereby providing a large, BRAF^V600E/K^ tumour-selective therapeutic window^3,4^. Attempts to overcome paradoxical RAF activation in RAS mutant cancer cells have included the development pan-RAFi that inhibit all RAF isoforms (ARAF, BRAF and CRAF)^19^ and ‘Paradox Breaker’ RAFis^20^. However, whilst these inhibitors show the desired effects on ERK1/2 activity in cells, they have not yet proved successful in clinical trials.

In the course of studying paradoxical RAF activation, we observed that many RAFi promoted a rapid (within 1 hour) inhibition of DNA replication. It is known that hyperactivation of ERK1/2 signalling beyond a critical ‘sweet spot’ can drive cell cycle arrest, senescence and even cell death ^21–23^; however, we found that the RAFi-induced inhibition of proliferation was independent of MEK1/2-ERK1/2 activation. Here we show that multiple RAFi, including all three clinically approved drugs (Vemurafenib, Dabrafenib and Encorafenib), pan-RAFi and Paradox Breaking RAFi induce rapid ERK1/2-independent activation of the Integrated Stress Response (ISR)^24^ to inhibit DNA synthesis. Mechanistically, RAFi drive ISR activation and cell cycle arrest by binding directly to GCN2 dimers and activating them in a manner reminiscent of the paradoxical activation of wild type RAF dimers by RAFi^13,18^. Since the ISR is a critical pathway that determines cell survival or cell death after stress^25^, these observations may be relevant to the further clinical optimisation of RAFi in the context of cancer cell stress responses.

## Results

### RAF inhibitors drive a rapid inhibition of DNA synthesis that is independent of ERK1/2 signalling

We analysed p-ERK1/2 and EdU incorporation using high-content microscopy^26,27^ in NCI-H358 cells (KRAS^G12C^-mutant lung adenocarcinoma cells) treated with Dabrafenib, a Group 1 RAFi (Fig 1A). Dabrafenib stimulated paradoxical activation of RAF, seen by the characteristic, broad bell-shaped concentration-response curve for p-ERK1/2. Given the adventitious tumour growth observed in the clinic^13,18^, we were surprised to see that Dabrafenib actually inhibited DNA synthesis, monitored by EdU incorporation (Fig 1A). The decline in EdU incorporation correlated with activation of ERK1/2 and was reversed at higher doses of Dabrafenib as p-ERK1/2 levels declined. We expanded our analysis to four additional RAFi (Encorafenib, Vemurafenib, PLX-4720 and GDC-0879), comparing them with cells treated with Trametinib (a MEK1/2 inhibitor or MEKi) (Supplementary Fig 1A & 1B). Trametinib caused a striking inhibition of p-ERK1/2 and EdU whereas all RAFi stimulated paradoxical activation of RAF and ERK1/2 signalling and four out of five RAFi also inhibited EdU incorporation (Supplementary Fig 1A & 1B). ERK1/2 signalling operates within a ‘sweet-spot’ to control cell proliferation, with high levels of p-ERK1/2 promoting cell cycle arrest^21–23^ so we initially attributed the inhibition of DNA synthesis to paradoxical RAF activation driving p-ERK1/2 beyond a level optimal for proliferation. However, close examination of the data argued against this. For example, Encorafenib was more potent than Dabrafenib at increasing p-ERK1/2 but less potent at inhibiting EdU incorporation, whilst GDC-0879 was equipotent to Dabrafenib at increasing p-ERK1/2 but completely failed to inhibit EdU incorporation (Supplementary Fig 1A and 1B). We therefore tested directly if ERK1/2 activation was required for RAFi-induced cell cycle arrest by combining Dabrafenib with Trametinib. Treatment with increasing doses of Trametinib caused a dose-dependent loss of both basal and paradoxical ERK1/2 activation (Fig 1B). The reduction in basal EdU incorporation caused by 5nM or 10nM Trametinib was partially reversed by Dabrafenib-dependent paradoxical ERK1/2 activation (Fig 1C), but Trametinib was unable to reverse the loss of EdU incorporation seen with 100nM-1µM Dabrafenib (Fig 1C). Identical results were obtained with the chemically distinct MEKi, Selumetinib (Supplementary Fig 1C and 1D). Together these results demonstrated that the RAFi-induced loss of EdU incorporation was independent of ERK1/2 activation and either reflected ERK1/2-independent signalling by RAF or an off-target effect of RAFi.

**Figure 1.**
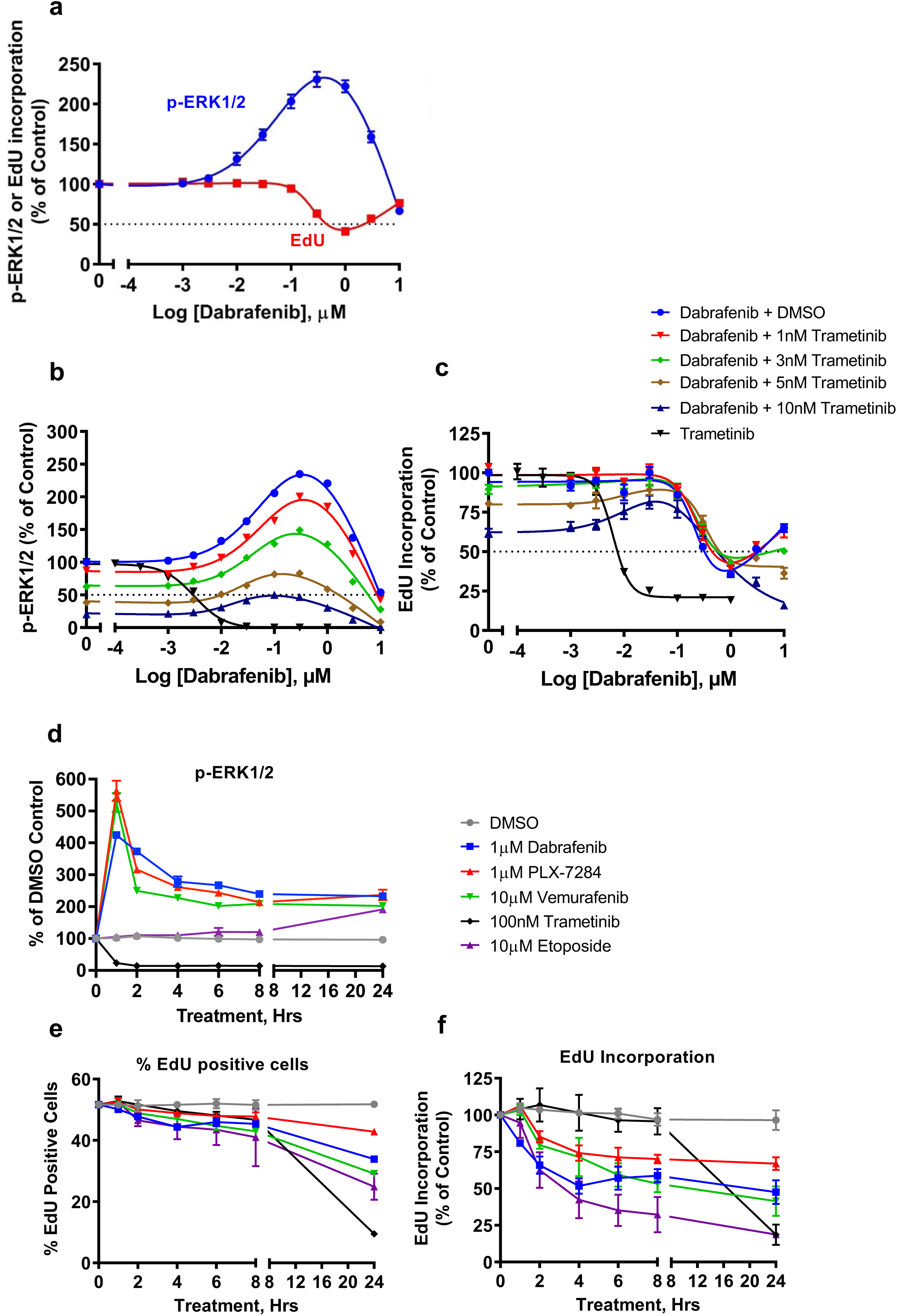
RAF inhibitors drive the rapid inhibition of DNA replication independent of paradoxical ERK1/2 activation. **a.** NCI-H358 cells treated with the indicated concentrations of Dabrafenib for 24 hours with a 10μM EdU pulse for the final hour. Cells were then fixed and permeabilized for EdU detection, immunofluorescence with a p-ERK1/2-specific antibody and co-stained with DAPI. Mean signal per cell was determined by high-content image analysis of 2000-15000 cells per condition. Normalised mean values ± SEM are shown, n = 3 replicate experiments **b & c.** NCI-H358 cells were treated with 1, 3, 5 or 10nM Trametinib for 1 hour prior to addition of the indicated concentration of Dabrafenib for 24 hours. Some cells received increasing concentrations of Trametinib alone for 24 hours. p-ERK1/2 (**b**) and EdU (**c**) incorporation were quantified by HCM analysis as above. Results shown are mean ± SEM of n=3 replicate experiments. **d-f.** NCI-H358 cells were treated with 1µM Dabrafenib, 1µM PLX-7284, 10µM Vemurafenib, 100nM Trametinib or 10µM Etoposide for the indicated times and analysed for p-ERK1/2 (**d**), percentage of EdU-postive cells (**e**) or intensity of EdU staining (**f**) as above. Results mean ± SEM of n=3 replicate experiments.

We examined the kinetics of RAFi-induced ERK1/2 activation and inhibition of EdU incorporation (Fig 1D-1F). Dabrafenib, Vemurafenib and PLX-7284 (a chemically distinct RAFi with properties similar to Vemurafenib and PLX-4720) caused rapid ERK1/2 activation that was maximal at 1 hour before declining into a sustained phase; Trametinib caused the immediate (1 hour) and sustained loss of p-ERK1/2. EdU staining revealed that RAFi-treated and MEKi-treated cells exhibited a comparable proportion of cells in S-phase for the first 8h of treatment (Fig 1E). However, within these ‘EdU-positive’ cells RAFi treatment caused a very rapid reduction in EdU staining intensity, consistent with inhibition of DNA replication (Fig 1F). This was apparent within 1h and reached a maximum 50% inhibition by 4h (Fig 1F) and was also seen by flow cytometry (Supplementary Fig 2A). This was not observed in MEKi-treated cells which exhibited a slower inhibition of DNA synthesis from between 8 and 24 hours. The rapid inhibition of DNA replication was reminiscent of an acute cell-cycle checkpoint such as the DNA damage response (DDR); indeed, the kinetics of the RAFi response matched that in cells treated with Etoposide (Fig 1F) but Dabrafenib failed to increase p-H2AX(S139) or p-CHK1(S345), both classic markers of DNA damage (Supplementary Fig 2B-2D). In summary, a variety of chemically distinct RAFi initiated a rapid inhibition of DNA replication that was independent of ERK1/2 activation, in a manner reminiscent of an acute DDR but without evidence of DNA damage.

### RAF inhibitors drive rapid, ERK1/2-independent activation of the unfolded protein response (UPR)

To identify RAFi-driven signalling pathways that were independent of MEK1/2-ERK1/2 signalling we performed an RNA-seq experiment. NCI-H358 cells were treated for 4 or 24 hours with 1µM Dabrafenib alone (Dab), 30nM Selumetinib (Sel) alone or the combination (Dab+Sel). These doses were previously optimised (Supplementary Fig 1C and 1D) and replicate samples with the same drug mastermix confirmed that Dab-driven activation of ERK1/2 was completely reversed by combination with 30nM Selumetinib (Dab+Sel) whilst Sel alone reduced basal p-ERK1/2 levels (Fig 2A). Analysis of the RNA-seq data confirmed that Sel reversed Dab-induced expression of *DUSP4*, *DUSP5* and *DUSP6* whilst Sel alone reduced the basal expression of these three well known ERK1/2 target genes (Supplementary Fig 3A); this showed that the experimental set-up for the RNA-seq had worked in identifying MEK1/2-dependent gene expression. Hallmark Genesets from the molecular signatures database were used to analyse the RNA-seq looking for gene sets induced or repressed by Dab and Dab+Sel (Supplementary Fig 3B). Dab-induced, MEK1/2-dependent gene sets such as ‘KRAS signalling Up’, ‘Epithelial Mesenchymal Transition’ and ‘Inflammatory responses’ also validated the experiment. Only three gene sets responded to Dab and Dab+Sel equally: ‘E2F targets’ and ‘G2M Checkpoint’ decreased strongly, likely reflecting the inhibition of DNA replication. The only gene set increased by RAFi in a MEK1/2-independent fashion was the ‘Unfolded Protein Response’. The UPR is a homeostatic signalling network activated by the accumulation of damaged or misfolded proteins in the ER lumen (ER stress) ^28,29^ and consists of three arms that are controlled by distinct stress sensors in the ER membrane. ATF6 translocates to the Golgi apparatus where it undergoes proteolytic cleavage before entering the nucleus to drive transcription of XBP-1 and components of the ER-associated degradation (ERAD) machinery. Inositol-requiring protein 1α (IRE1α) coordinates with ATF6 activation by driving the splicing of mRNA encoding the XBP-1 transcription factor; removal of an intron results in expression of the active XBP-1s transcription factor which drives expression of ER chaperones. Finally, Protein kinase RNA-like ER kinase (PERK) phosphorylates translation initiation factor eIF2α; this attenuates protein synthesis but allows non-canonical translation of ATF4, which in turn drives expression of CHOP and ATF3. Further analysis of the RNA-seq data confirmed strong Dab-induced, MEK1/2-independent expression of 10 genes that are well known markers of the UPR (Supplementary Fig 3C) including ATF4, ATF3, CEBPB, DDIT3 (CHOP), DDIT4 (REDD1) and SESN2. When we examined the RNA-seq replicate cell extracts (Fig 2A) we observed a striking increase in expression of ATF4, CHOP and their downstream target TRIB3, an ER-sensing pseudokinase^30^, following 4 or 24 hours Dab treatment; this was unaffected by Sel and was therefore independent of MEK1/2-ERK1/2 (Fig 2A).

**Figure 2.**
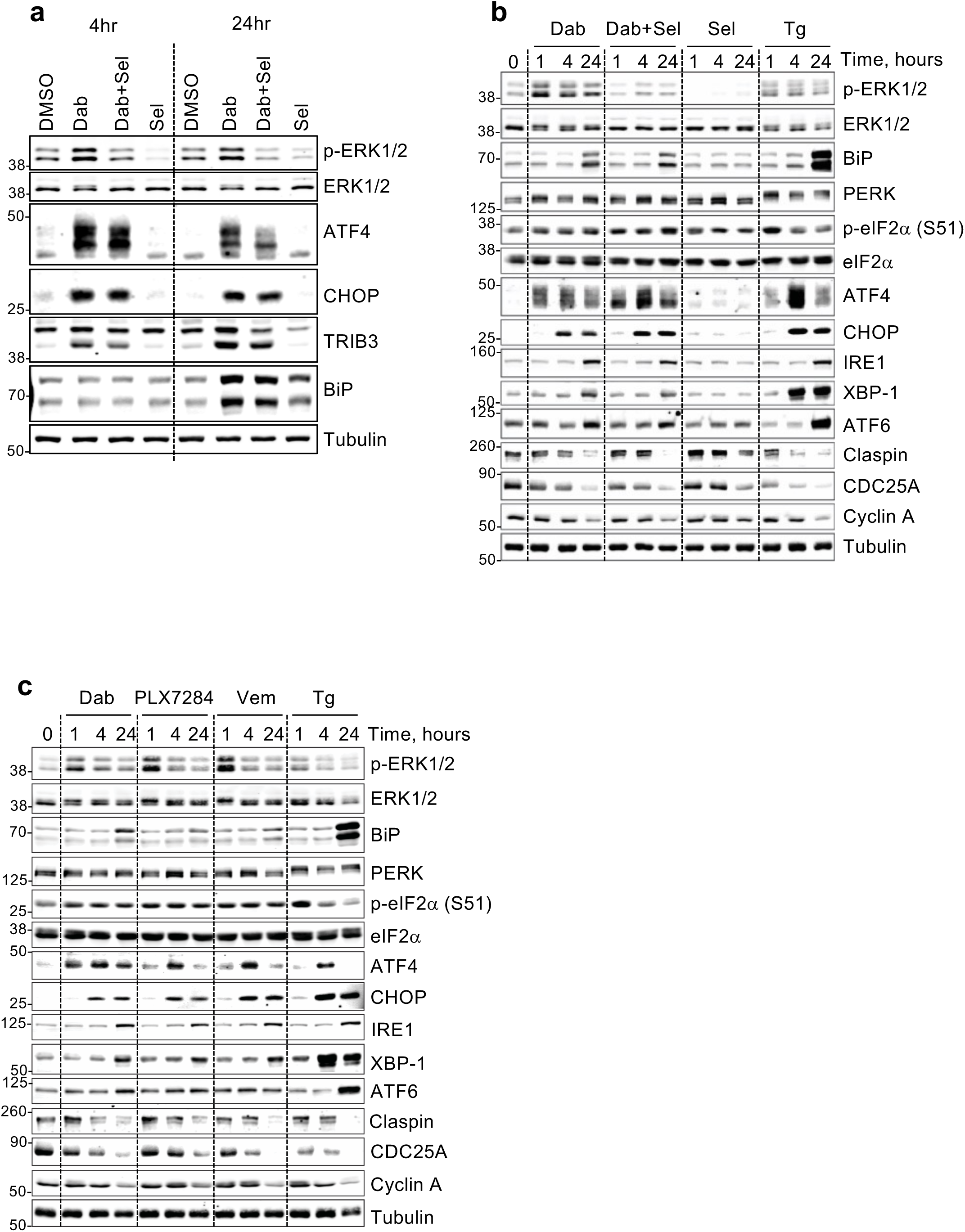
RAFi drive rapid, ERK1/2-independent expression of ATF4 and CHOP but not other arms of the UPR. **a.** NCI-H358 cells were treated with 1µM Dabrafenib (Dab), 30nM Selumetinib (Sel) or the combination (Dab+Sel) for 4 or 24 hours. Samples were processed for RNA-seq (Supplementary Figure 3) and replicates dishes were used to prepare whole cell extracts that were resolved by SDS-PAGE and immunoblotted with the indicated antibodies. **b.** NCI-H358 cells were treated with Dab, Sel and Dab+Sel as above and additionally with 100nM Thapsigargin for 1, 4 or 24 hours. Whole cell lysates were fractionated by SDS-PAGE and immunoblotted with the indicated antibodies. **c.** NCI-H358 cells were treated with 1µM Dabrafenib (Dab), 1 µM PLX-7284 (PLX), 10µM Vemurafenib (Vem), or 100nM Thapsigargin (Tg) for 1, 4, or 24 hours. Whole cell extracts were resolved by SDS-PAGE and immunoblotted with the indicated antibodies. All results are representative of three independent experiments and in all cases results were captured by LiCor.

To investigate the UPR in more detail, NCI-H358 cells were treated with Thapsigargin (Tg), an ER stress agonist, which activated of PERK (indicated by its hyperphosphorylation) and increased expression of ATF4, CHOP, XBP-1 and ATF6 consistent with activation of all arms of the UPR (Fig 2B). Dabrafenib stimulated a rapid (1 hour) increase in ATF4 and a slower increase in CHOP but little activation of PERK and only modest expression of BIP, ATF6 or XBP1. Selumetinib alone had no effect on ER stress markers but effectively inhibited basal and Dab-induced p-ERK1/2. Critically, Selumetinib did not reverse the Dab-induced expression of ATF4 or CHOP. A similar profile of ATF4 and CHOP expression but modest and delayed expression of other markers of ER stress was seen with two other RAFi, PLX-7284 and Vemurafenib (Fig 2C). To further validate this finding, we established a HCM assay for ATF4, CHOP and XBP-1 using Tg or Tunicamycin (Tn) as positive controls (Supplementary Figure 4A-4C). This demonstrated that both RAFi, Dabrafenib and PLX-7284, induced ATF4 expression with similar kinetics and to the same extent as Tg and Tn. However, whilst Dabrafenib and PLX-7284 increased CHOP and XBP-1 expression, the magnitude of this response was far less than that seen with Tg or Tn; CHOP and XBP-1 expression was much more responsive to ER stressors.

To test the involvement of PERK we used the potent and selective PERK inhibitor GSK2606414^31^. This blocked the hyperphosphorylation of PERK, the expression of ATF4, CHOP and the slower expression of ATF6 in response to Thapsigargin, but had no effect on the expression of ATF4 or CHOP in response to Dabrafenib (Fig 3A). We also used HCM to quantify the effects of the PERK inhibitor; GSK2606414 inhibited CHOP expression (Fig 3B) and reversed the inhibition of EdU incorporation (Fig 3C) induced by Tg or Tn in a concentration-dependent fashion, indicating that PERK was required for these responses to ER stress. In contrast, the inhibition of EdU incorporation induced by Dabrafenib, Vemurafenib or PLX-7284 was unaffected by GSK2606414 (Fig 3D). In summary, these data showed that RAFi drive the rapid expression of ATF4 to the same magnitude as the ER stressors, Tg and Tn, but cause little or no activation of the other arms of the UPR. Furthermore, PERK plays no role in the RAFi-induced expression of ATF4 or CHOP or RAFi-induced inhibition of cell proliferation.

**Figure 3.**
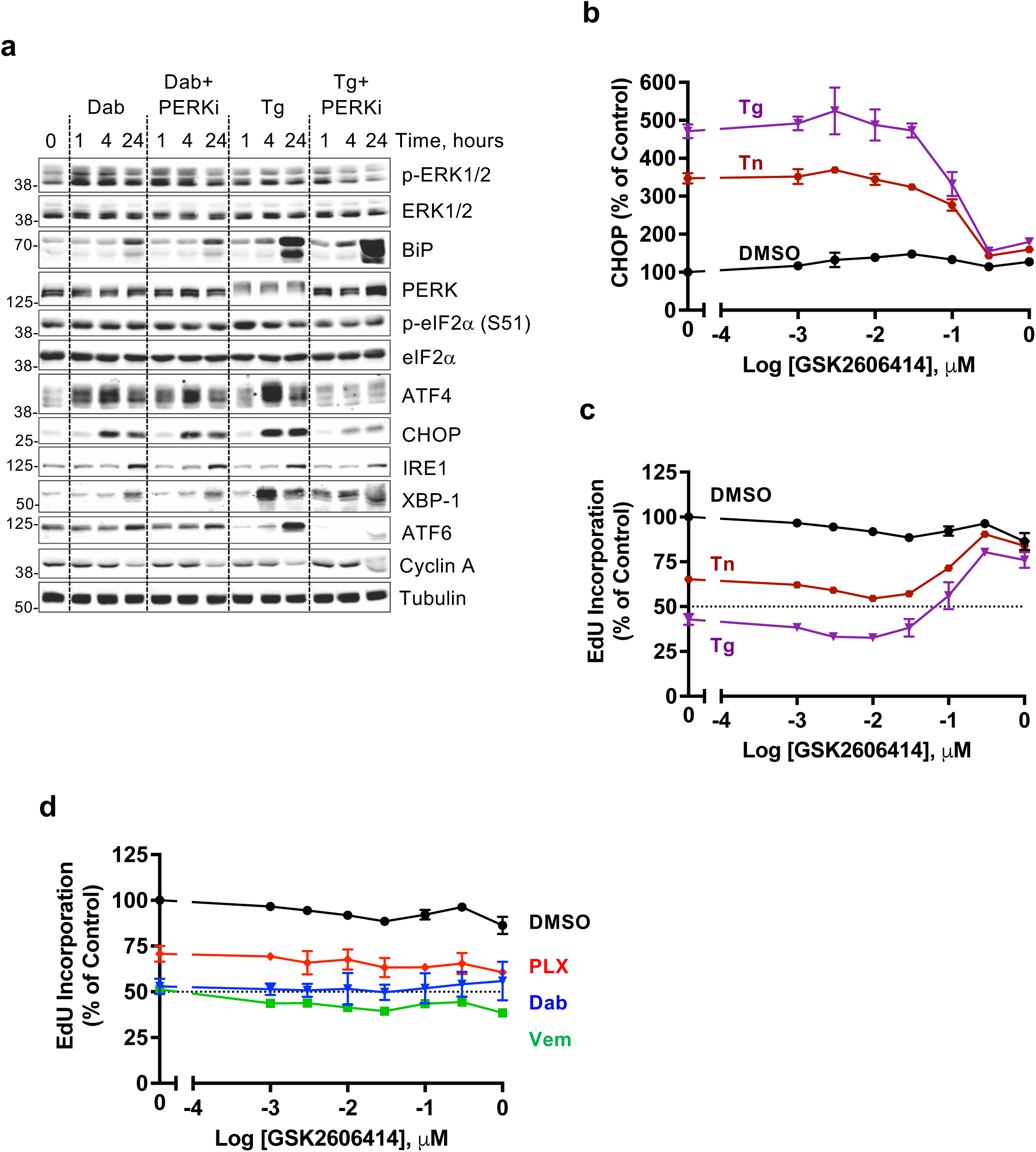
PERK plays no role in the RAFi-induced expression of ATF4 or CHOP or inhibition of cell proliferation. **a.** NCI-H358 cells were pretreated for 30 minutes with either DMSO or 300nM GSK2606414 (PERK inhibitor) prior to the addition of 1µM Dabrafenib or 100nM Thapsigargin for 1, 4 or 24 hours. Whole cell extracts were resolved by SDS-PAGE and immunoblots with the indicated antibodies were captured by LiCor. Results are representative of three independent experiments. **b. & c.** High content microscopy (HCM) analysis of CHOP expression and EdU incorporation in NCI-H358 cells treated with the indicated concentrations of GSK2606414 for 30 mins prior to the addition of DMSO, 100nM Thapsigargin (Tg) or 2µg/ml Tunicamycin (Tn) for 24 hours. Cells were analysed for EdU incorporation as in Figure 1. Mean signal per cell was quantified by high-content image analysis of 2000-15000 cells per condition. **d.** NCI-H358 cells were treated with the indicated concentrations of the GSK2606414 for 30 mins before the addition of 1µM Dabrafenib, 1 µM PLX-7284 or 10µM Vemurafenib for 24 hours. Cells were analysed for EdU incorporation as above. For **b-d** normalised mean values ± SEM are shown from n = 3 replicate experiments.

### RAF inhibitors activate ISR and inhibit cell proliferation by activating GCN2

The phosphorylation of eIF2α attenuates protein synthesis but allows non-canonical translation of ATF4 to drive a homeostatic gene expression programme, allowing cells to adapt to various forms of stress; this ancient signalling pathway is called the Integrated Stress Response (ISR)^24^. The rapid expression of ATF4 suggested that RAFi were able to activate the ISR, but this was apparently not dependent on PERK. Three other eIF2α kinases (EIF2AKs) can phosphorylate Ser51 of eIF2α to initiate the ISR: PKR (Double-stranded RNA-dependent protein kinase) is activated by viral infection, HRI (Heme-regulated eIF2α kinase) by Haem depletion and GCN2 (General control non-depressible protein 2) by amino acid starvation. We examined a potential role for GCN2 since an antibody that detects the activating phosphorylation site at Thr899 was available. Four different RAFi (Dabrafenib, PLX-7284, Vemurafenib and LY-3009120) increased p-T899 GCN2 and p-S51 eIF2α, increased the expression of ATF4, CHOP and TRIB3 (markers of the ISR) and caused a loss of CDC25A (which is linked to inhibition of DNA replication) (Fig 4A).

**Figure 4.**
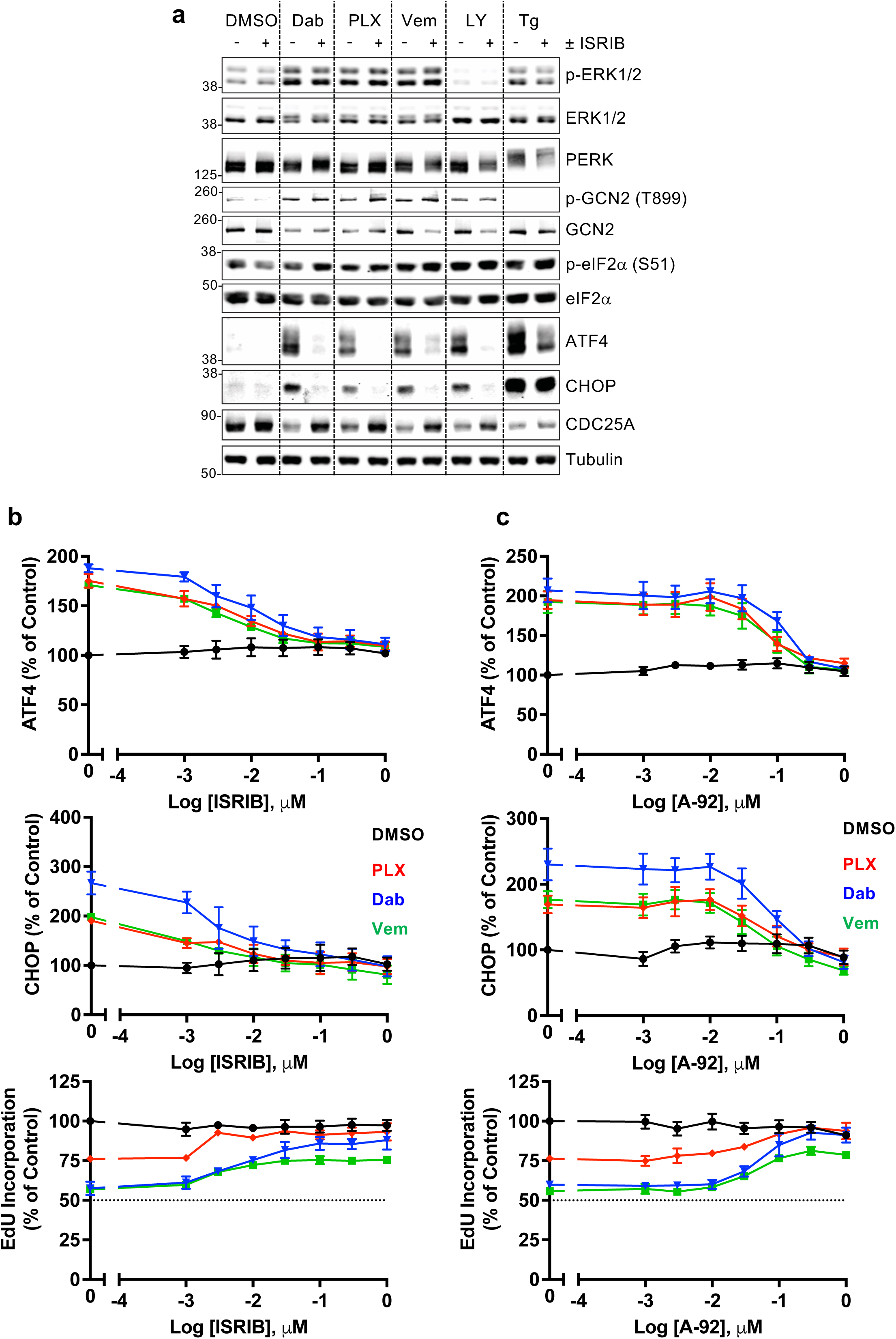
RAF inhibitors activate ISR and inhibit cell proliferation by activating GCN2. **a.** NCI-H358 cells were treated with DMSO or 300nM ISRIB for 30 mins prior to the addition of 1µM Dabrafenib (Dab), 1 µM PLX-7284 (PLX), 10µM Vemurafenib (Vem), 1 µM LY-3009120 (LY) or 100nM Thapsigargin (Tg) for 4 hours. Whole cell lysates were fractionated by SDS-PAGE and immunoblotted with the indicated antibodies. Western blots from a single experiment representative of 3 experiments are shown**. b. & c.** NCI-H358 cells were treated with the indicated concentrations of either ISRIB (**b**) or A-92 (**c**) for 30 mins prior to the addition of DMSO, 1µM Dabrafenib (Dab), 1 µM PLX-7284 (PLX) or 10µM Vemurafenib (Vem). Cells were treated for 7 hours with a pulse of 10μM EdU for the final hour, then fixed and permeabilized for EdU detection, immunofluorescence with ATF4- or CHOP-specific antibodies and co-stained with DAPI. Mean signal per cell was quantified by high-content image analysis of 2000-15000 cells per condition. Normalised mean values ± SEM are shown, from n = 3 replicate experiments.

Thapsigargin failed to activate GCN2 but caused hyperphosphorylation of PERK, activation of the ISR and loss of CDC25A (Fig 4A). The pan-RAFi LY-3009120 inhibits all RAF isoforms so fails to drive paradoxical ERK1/2 activation but still activated GCN2 corroborating that RAFi-dependent activation of GCN2 and the ISR was independent of ERK1/2 activation (Fig 4A). The eIF2α antagonist ISRIB^32,33^ completely reversed the RAFi-induced expression of ATF4, CHOP and TRIB3; it also reversed the loss of CDC25A, indicating that this was a consequence of the ISR (Fig 4A). Similar results were observed with two further RAFi, Encorafenib and PLX-8394 (Plixorafenib) (Supplementary Fig 5A). We also assessed ATF4 and CHOP expression and EdU incorporation by HCM. ISRIB inhibited the expression of ATF4 and CHOP and reversed the inhibition of EdU incorporation induced by Dabrafenib, PLX-7284 and Vemurafenib (Fig 4B) but had little if any effect on the expression of ATF4, CHOP or loss of EdU incorporation seen with Tg and Tn (Supplementary Fig 5B) suggesting that these ER stressors exert their effects via another arm of the UPR. Finally, A-92^34^, an ATP-competitive GCN2 kinase inhibitor reversed the expression of ATF4 and CHOP and the inhibition of EdU incorporation induced by Dabrafenib, PLX7284 and Vemurafenib (Fig 4C) and similar results were seen for PLX-8394, LY-3009120 and Encorafenib (Supplementary Fig 4A). In contrast, A-92 had no effect on the response to the ER stressors Tg and Tn (Supplementary Fig 5C).

We also employed genetic knockdown or knockout of GCN2 and expanded our analysis into different cell types. siRNA-mediated knock-down of GCN2 strongly inhibited Dab-driven expression of ATF4 and CHOP in NCI-H358 cells, whilst siRNA against ATF4 strongly inhibited Dab-driven CHOP expression without affecting GCN2 abundance (Fig 5A). Dab increased expression of ATF4 in wild type immortalised MEFs (iMEFs) and PERK KO iMEFs, but this response was greatly reduced in GCN2 knockout iMEFs (Fig 5B). Attempts to generate GCN2 KO NCI-H358 cells using CRISPR/Cas9 gene editing were unsuccessful; however, knockout of GCN2 was successful in HCT116 colorectal cancer cells (Fig 5C & 5D). In these cells GCN2 KO abolished expression of ATF4 and CHOP induced by Dab and Histidinol, an inhibitor of histidyl tRNA synthetase that mimics histidine starvation, thereby activating GCN2 through accumulation of uncharged tRNAs (Fig 5C); in contrast, the response to the ER stressor Tunicamycin was unaffected by GCN2 KO. Interestingly a separate GCN2 inhibitor (GCN2iB) caused a GCN2-dependent increase in ATF4 and CHOP expression (Fig 5C). We expanded this analysis by comparing four different RAFi: Dabrafenib, LY-3009120 and Encorafenib all increased p-T899 GCN2, p-S51 eIF2α and ATF4 and CHOP expression and these responses were almost completely GCN2-dependent (Fig 5D); in contrast GDC-0879 failed to activate GCN2 or the ISR (Fig 5D). Finally, we used HCM assays for p-ERK1/2 and ATF4 to screen additional RAFi using comparisons with the previous inhibitors (Supplementary Fig 6A-D). Paradox breaker RAFis (PLX-8394 and PLX-7922) derived from Vemurafenib^20^ were less potent and effective for ERK1/2 activation but still increased ATF4 expression. Both pan-RAF inhibitors, LY-3009120 and AZ628, strongly inhibited ERK1/2 activation. LY-3009120 increased ATF4 expression over the same concentration range as it inhibited p-ERK1/2; in contrast, AZ628 was very poor at increasing ATF4 despite strong p-ERK1/2 inhibition.

**Figure 5.**
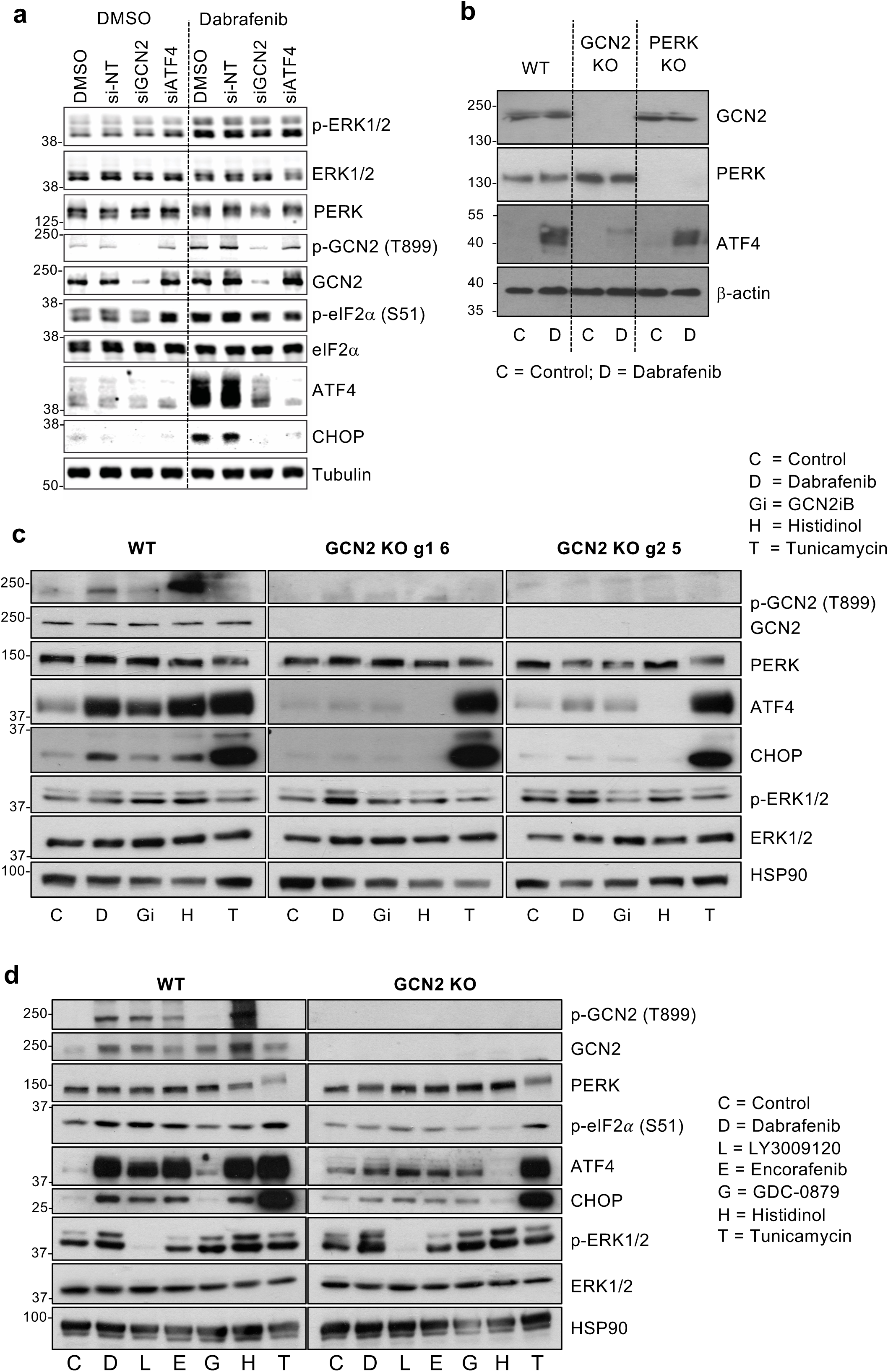
Knockdown or knockout of GCN2 blocks RAFi-induced activation of the ISR. **a.** NCI-H358 cells were either mock transfected or transfected with 10nM of siRNA either non targeting (siN-T) or targeted to GCN2 (siGCN2) or ATF4 (siATF4) for 48 hours cells before treatment with DMSO or 1µM Dabrafenib for 4 hours. **b.** Immortalised wild type, GCN2 KO or PERK KO mouse embryonic fibroblasts were treated with DMSO (C, control) or 1 µM Dabrafenib (D) for 4 hours. **c.** Wildtype HCT116 or 2 independent CRISPR generated GCN2 knockout (GCN2KO) clones; guide 1 clone 6 (g1 6) or guide 2 clone 5 (g2 5) were treated with 1µM Dabrafenib (D), 0.03 µM GCN2iB, 16 mM Histidinol or 200nM Tunicamycin (T) for 4 hours. **d.** Wild type or GCN2 KO (clone g1 6) HCT116 cells were treated with either DMSO (C, control), 1µM Dabrafenib (D), 1 µM LY-3009120, 3 µM Encorafenib, 1 µM GDC-8079 (G), 16mM Histidinol (H) or 200nM Tunicamycin (T) for 4 hours. For all experiments (**a-d**) whole cell lysates were fractionated by SDS-PAGE and immunoblotted with the indicated antibodies. Results were captured by LiCor (**a**) or conventional ECL (**b-d**) from a single experiment representative of n=3 for all experiments (**a-d**).

Taken together, these results using small molecule inhibitors (ISRIB and the GCN2 inhibitor A-92) and genetic knockdown or knockout of GCN2 demonstrated unequivocally that 8 of 10 RAFi studied activated the ISR in cells and this required GCN2 kinase activity. Only GDC-0879 and AZ628 failed to activate GCN2 and/or the ISR.

### RAFi bind directly to GCN2 dimers and drive their activation *in vitro* and in cells

In considering how RAFi might activate GCN2 we noted that Dabrafenib rapidly increased p-T899 GCN2 and p-S51 eIF2α (15-30 mins) and ATF4 expression (30min-1hour) (Fig 6A); the increase in ATF4 expression was notably more rapid than that seen with Thapsigargin, a validated ISR agonist. Similar, rapid phosphorylation of GCN2 and expression of ATF4 was observed in Dabrafenib-stimulated colorectal cancer cells, regardless of whether they exhibited paradoxical activation of ERK1/2 (HCT116, cells KRAS^G13D^) or inhibition of ERK1/2 (HT29 cells, BRAF^V600E^) (Fig 6B). HT29 cells had much lower expression of GCN2 than HCT116 cells and exhibited lower expression of ATF4 and CHOP, but both cells responded to Dabrafenib with a rapid phosphorylation of GCN2. The rapid activation of GCN2 and ATF4 expression was reminiscent of ERK1/2 activation (apparent at 15-30 mins) arising from Dabrafenib-induced activation of wild type RAF dimers (Fig 1D and 6A) so we tested the possibility that GCN2 activation might be a RAF-dependent response; for example, RAF activation by RAFi might allow RAF to phosphorylate GCN2 or interact with GCN2 to facilitate its activation. However, in western blot analysis siRNA-mediated knockdown of CRAF alone prevented Dabrafenib-induced ERK1/2 activation but not Dabrafenib-induced phosphorylation of p-T899 GCN2, p-S51 eIF2α or expression of ATF4 and CHOP (Supplementary Fig 7A). Similar results were obtained in high content microscopy analysis, where CRAF knockdown prevented ERK1/2 activation by six different RAFi but had no effect on RAFi-induced ATF4 expression (Supplementary Fig 7B-E). We considered the possibility that RAFi might disrupt amino acid metabolism, mimicking amino acid starvation, but the kinetics of GCN2-ISR activation seemed too rapid to support this possibility.

**Figure 6.**
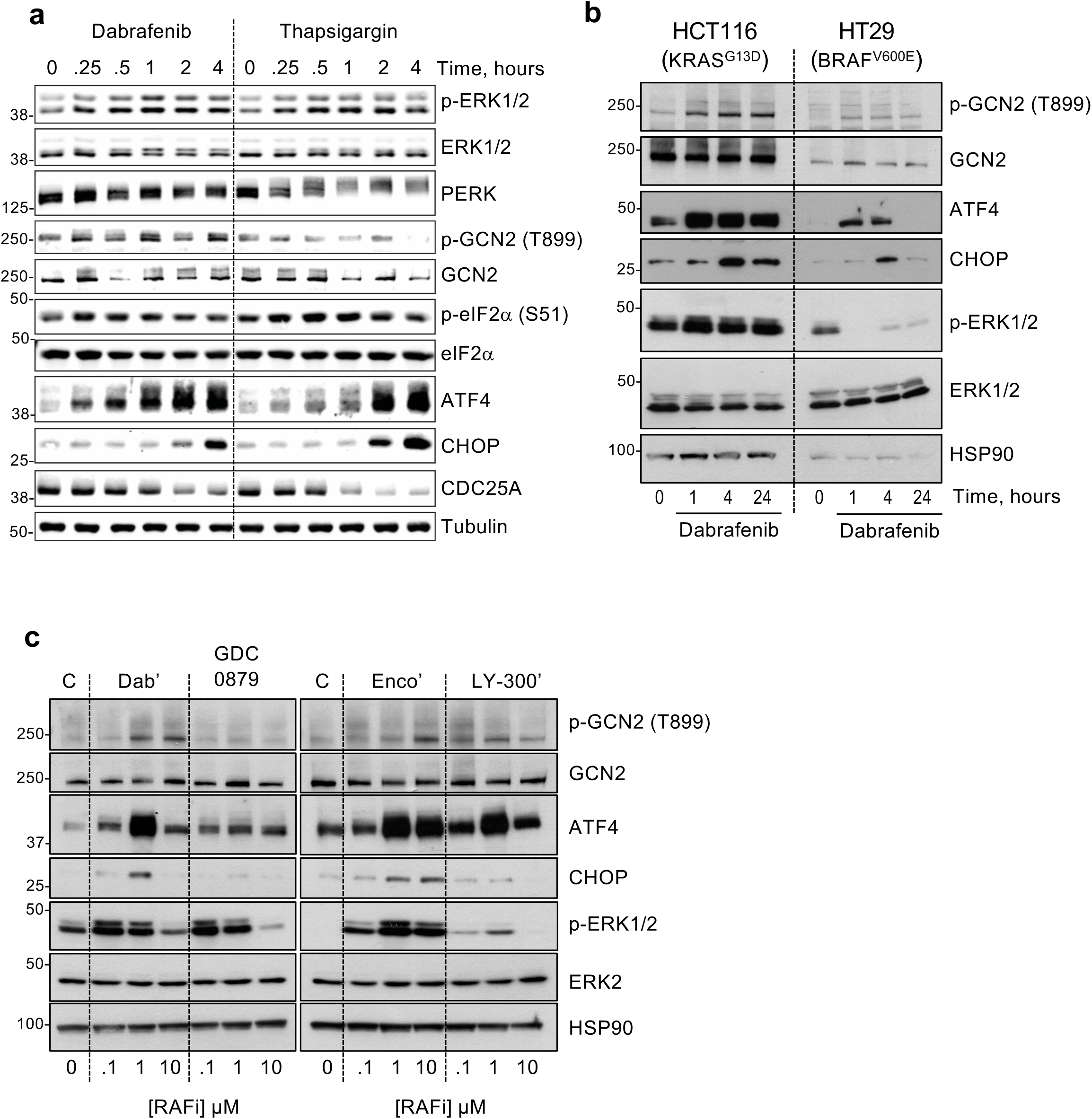
RAF inhibitors activate GCN2 and the ISR rapidly with a bell-shaped concentration response curve. **a.** NCI-H358 cells were treated with 1µM Dabrafenib or 200nM thapsigargin for up to 4 hours. Whole cell extracts were resolved by SDS-PAGE and immunoblotted with the indicated antibodies. **b.** HCT116 or HT29 cells were treated with 1µM Dabrafenib for up to 4 hours and whole cell lysates were analysed as above. **c.** HCT116 cells were treated with either DMSO (C, control) or the indicated concentrations of Dabrafenib (Dab’), GDC-8079 (GDC), Encorafenib (Enco’) or LY-3009120 (LY-300’) and western blots performed as in **a**. Results were captured by LiCor (**a**) or conventional ECL (**b, c**) from a single experiment representative of n=3 for all experiments (**a-c**).

We performed concentration-response curves with four different RAFi in HCT116 cells. Dabrafenib, Encorafenib and LY-3002190 all activated GCN2 and ATF4 expression whereas GDC-0879 again did not (Fig 6C). Interestingly, ATF4 expression (a strong, dynamic readout of ISR activation) increased at 100nM Dabrafenib and peaked at 1µM before declining at 10µM. Similar ‘bell-shaped’ concentration-response curves were observed for Encorafenib and LY-3002190 and were reminiscent of ERK1/2 activation arising from RAFi-induced activation of RAF dimers (Fig 1A; Supplementary Fig 1A). Previous studies showed that some early stage RAFi exhibit high affinity for the GCN2 kinase domain ^35–37^. Furthermore, crystal structures of human GCN2 exhibit a back-to-back dimer configuration ^37,38^ consistent with the active dimer conformations of other protein kinases including PKR, IRE1 and BRAF ^13,39^. Dimerization is observed during ‘normal’ RAS-dependent RAF activation and is critical for paradoxical activation of RAF by RAFi ^13,18^. Similarly, dimerization is critical for activation of the GCN2 kinase domain ^38, 40, 41^.

Prompted by our results and these precedents we considered the hypothesis that many RAFi are also GCN2 inhibitors that promote paradoxical activation of GCN2 dimers; to test this we first assessed the effect RAFi on GCN2 *in vitro*. When purified, recombinant, full-length human GCN2 was incubated with increasing doses of Dabrafenib in *in vitro* kinase assays we observed a concentration-dependent increase in p-T899 indicating activation and autophosphorylation of GCN2 (Fig 7A); furthermore, as with the cellular response (Fig 6C), this response was also ‘bell-shaped’, peaking at 0.5-1µM before declining at higher doses (Fig 7A). In contrast, GDC-0879 failed to increase p-T899 GCN2 at any doses (Fig 7B), agreeing with its failure to activate GCN2 in cells (Fig 5D and 6C). Smaller concentration response curves to allow side-by-side comparison of four RAFi demonstrated that LY-3009120, Encorafenib and Dabrafenib activated purified GCN2 at 100nM, peaking at 1µM and then declining at 10µM whereas GDC-0879 again failed to activate GCN2 (Fig 7C). In Fig 7C we also included purified eIF2α as a substrate for GCN2 and observed that p-S51 eIF2α increased slightly over the same range of doses as p-T899 with LY-3009120, Encorafenib and Dabrafenib. However, this was a very modest effect compared to the increase in p-S51 eIF2α observed when excess ATP was added to the *in vitro* kinase assays to activate GCN2. In summary, these results demonstrated that LY-3009120, Encorafenib and Dabrafenib could both bind to, and activate, purified GCN2 *in vitro*, mirroring their effects in cells and confirming a RAF-independent property for these RAFis at doses at which they activate ERK1/2 signalling.

**Figure 7.**
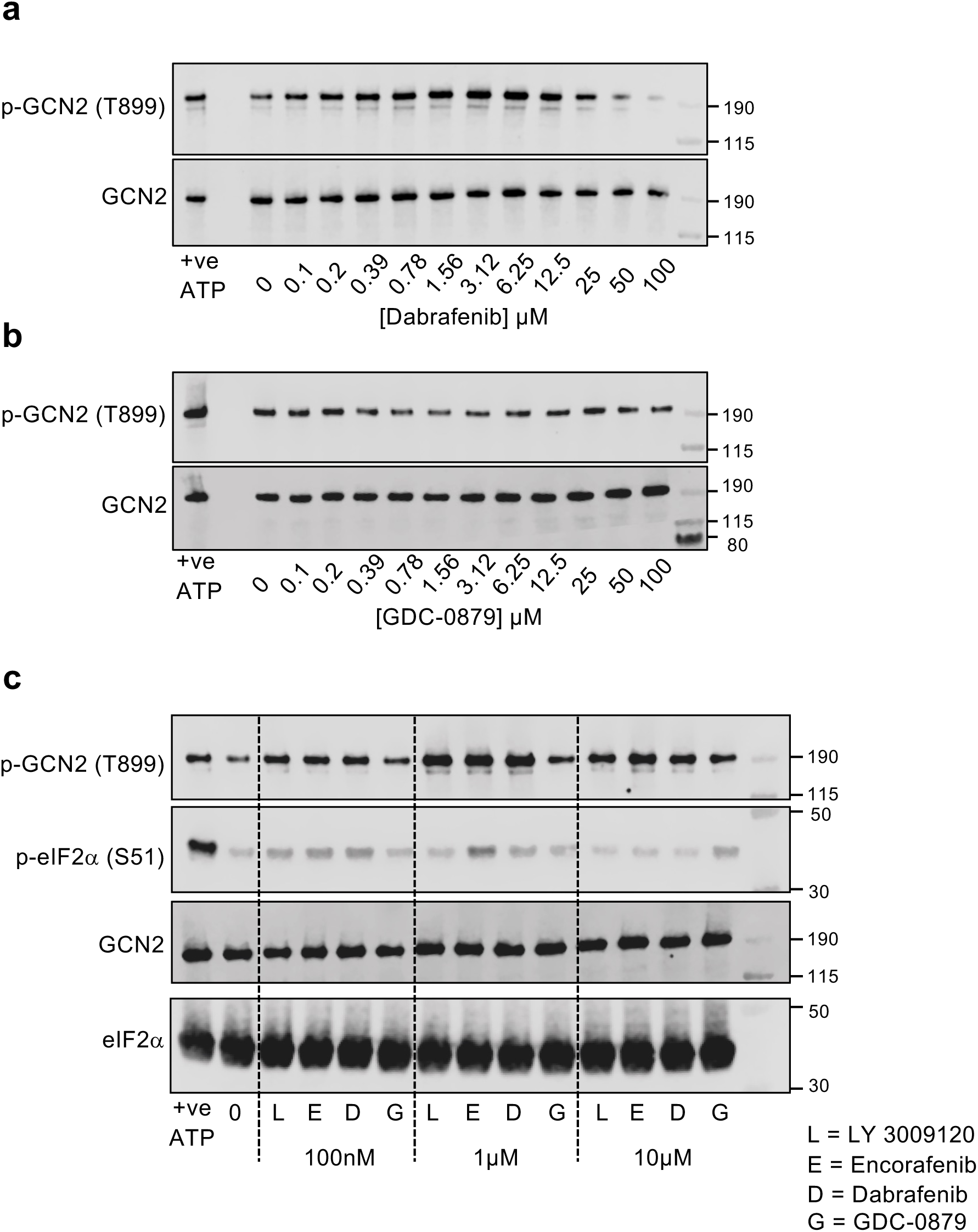
RAF inhibitors bind directly and activate GCN2 *in vitro* with a bell-shaped concentration response curve. a & b. 10 nM purified recombinant full length human GCN2 was incubated at 30°C for 20 mins with 10 µM ATP and the indicated concentrations of Dabrafenib (**a**) or GDC-8079 (**b**). Samples were resolved by SDS-PAGE and probed with the indicated antibodies. **c.** Reactions were performed as above with 100nM, 1 µM or 10µM of either LY-3009120 (L), Encorafenib (E), Dabrafenib (D) or GDC-8079 (G) and also included 2 µM eIF2α. Samples were resolved by SDS-PAGE and immunoblotted with the indicated antibodies. Single experiments representative of n = 3 are shown.

### Structural and mutational insights into RAFi-dependent activation of GCN2

Given the ability of RAFi to bind and activate GCN2, we generated structural models of GCN2-RAFi complexes. In common with BRAF, crystal structures of active human GCN2 exhibit a back-to-back dimer conformation ^37,38^ (Fig 8A). Cross-dimer interactions between the side chains of Tyr651 and Tyr652 are central in the dimeric interface (Fig 8B) and are dependent on Tyr651 occupying a ‘Tyr-up’ position. In contrast, these interactions are not observed in the alternative dimeric arrangement of yeast GCN2, where the side chain of Tyr651 points towards the kinase active site ^42^. This ‘Tyr-down’ configuration blocks catalytic output in other kinases (Nek7 and IRE1) and its reversal allows back-to-back dimerization and kinase activation^43^. In an active kinase, the Tyr651 side chain forms the top R4 position in the regulatory spine (R-spine), a hallmark of active kinase structures, which comprises four hydrophobic side chains stacked in four specific positions, termed RS1-RS4, (Fig 8B) ^44,45^. Critically, the R-spine cannot assemble when Tyr651 is in a Tyr-down position because the Tyr651 side chain occupies the RS3 space, leaving RS4 empty. Dabrafenib, a Type I.5 inhibitor, is predicted to bind to GCN2 and occupy the RS3 space; this is only compatible with a Tyr-up position, thereby stabilising an active conformation that promotes an active back-to-back dimer (Fig 8C). At low concentrations of Dabrafenib this would inhibit one protomer but paradoxically stimulate the kinase activity of the second, drug-free, ATP-bound protomer in the dimer. At higher concentrations of Dabrafenib, both kinase sites would be occupied and inhibited. In the model of GCN2 bound to LY3002190, a Type II inhibitor, the RS3 space is occupied by Leu640 and LY3002190 promotes a DFG-out, C-helix in, Tyr-up conformation (Fig 8D), thereby assembling the R-spine to promote an active back-to-back dimer which supports paradoxical activation. In contrast, GDC-0879, a Type I inhibitor, does not occupy the RS3 space so the R-spine does not assemble. GCN2 has a bulkier gatekeeper residue (Met) compared with BRAF (Thr), and GDC-0879 binding is predicted to dock in an orientation that fails to stabilise an active conformation (Fig 8E).

**Figure 8.**
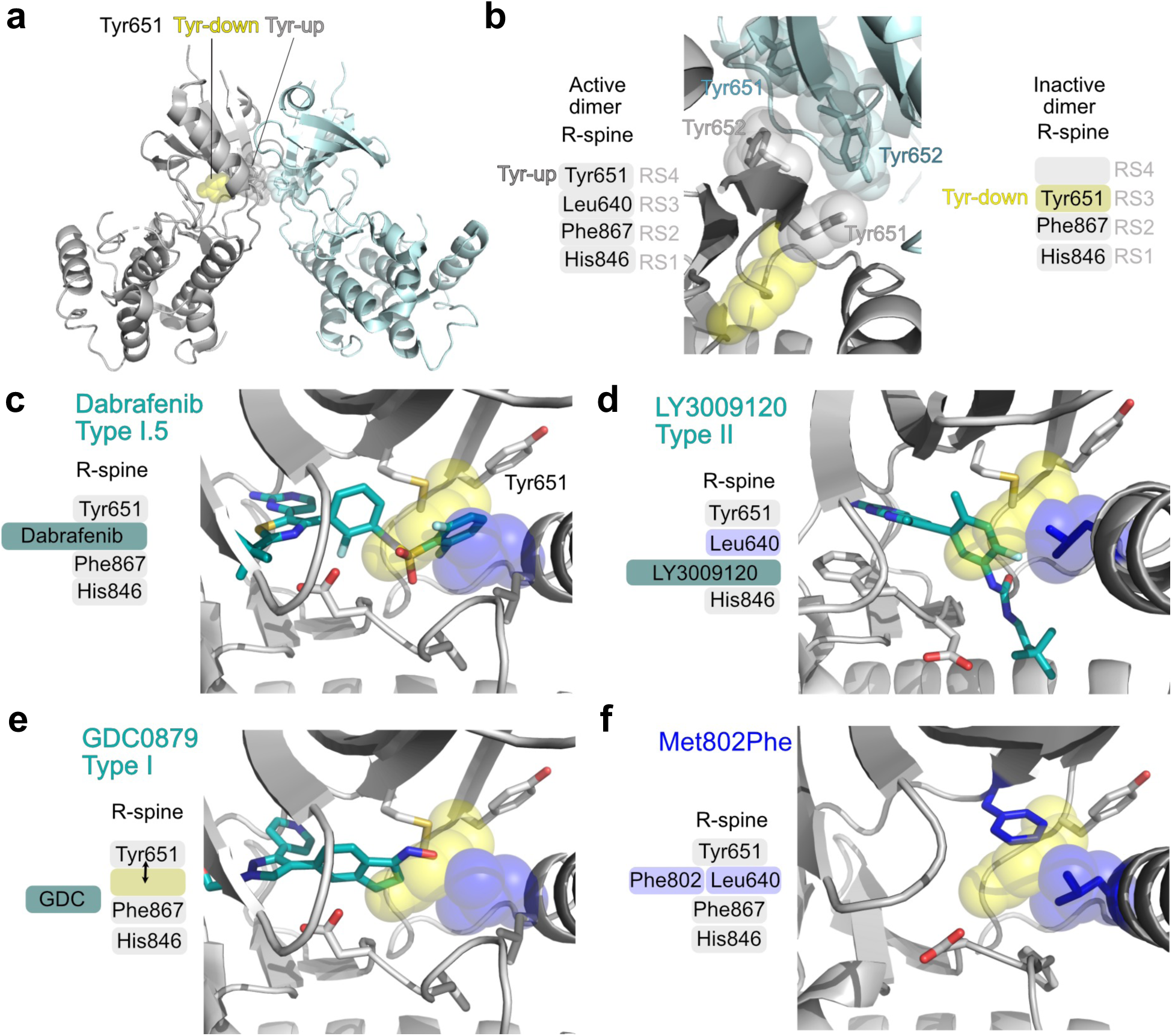
Structural models of RAF inhibitors binding to GCN2. Structural models of GCN2-ligand complexes were generated to provide a rationale for their biochemical properties. **a.** Crystal structures of human GCN2 exhibit a back-to-back dimer, consistent with the active dimer conformations of PKR, IRE1 and other kinases. **b.** Interactions between the side chains of Tyr651 (grey) and Tyr652 (blue) are central in the dimeric interface. The Tyr side chain forms the top position in the regulatory spine (R-spine), a hallmark of active kinase structures comprising four hydrophobic side chains that occupy specific positions (RS1-RS4). The R-spine cannot assemble when Tyr651 is in a Tyr-down position because the Tyr side chain occupies the space of RS3 and RS4 is empty. The Tyr-down position and RS3 space are shown as yellow and blue space filling spheres, respectively. **c.** Dabrafenib binding to GCN2 is predicted to occupy the RS3 space and Tyr-651 is in the up position **d.** The structural modelling of LY3002190 is consistent with a DFG-out, C-helix in, Tyr-up conformation with Leu640 occupying the RS3 space. **e.** GDC0879, a Type I inhibitor, does not protrude into the RS3 space, which is left unoccupied; this orientation is not predicted to stabilise an active conformation. **f.** Mutation of Met802 to a bulkier Phe residue is predicted to have a substantial clash with the Tyr-down position, stabilising Tyr-up through direct interactions with Leu640 in the RS3 space. See text for details.

These structural predictions are entirely consistent with the results we observed for RAFi activation of GCN2 and the ISR in cells but were predicated on the assumption that the RAFi bind to the GCN2 canonical kinase domain to elicit its activation. However, full length GCN2 contains both a conventional kinase domain and a pseudokinase domain^46^ which could feasibly support RAFi binding and drive allosteric (kinase-independent) signalling through conformational changes in signalling complexes. Since our results with the conventional GCN2 kinase inhibitor A-92 suggested a requirement for the canonical kinase domain, we generated two different catalytically inactive (kinase dead, or KD) mutants EGFP-GCN2^K619A^ or EGFP-GCN2^D848N^ and transiently expressed them in GCN2 KO HCT116 cells, comparing outputs with wild type EGFP-GCN2. WT GCN2 restored Dabrafenib-induced autophosphorylation of GCN2 at T899, whereas the GCN2 KD mutants failed to do so (Fig 9A). Protein kinases typically have a ‘gatekeeper’ residue whose molecular identity can dictate the binding affinity of ATP-competitive kinase inhibitors^47^. Mutation of this site can influence inhibitor binding, basal kinase activity and confer drug resistance^12, 48,49^. In BRAF the gatekeeper residue is Thr529 whereas in GCN2 it is the bulky Met802. We generated and transiently expressed four different GCN2 gatekeeper mutants in GCN2 KO HCT116 cells, using WT GCN2 as a control. Both EGFP-GCN2^M802A^ and EGFP-GCN2^M802G^ exhibited little or no activation in response to Dabrafenib indicating that they acted as drug-resistant mutants (Fig 9A); this confirmed that Dabrafenib was binding to the canonical kinase domain to activate GCN2, rather than the pseudokinase domain. Interestingly, the even bulkier gatekeeper mutant, EGFP-GCN2^M802F^ acted as a constitutively active kinase, exhibiting strong p-T899 even in the absence of Dabrafenib. EGFP-GCN2^M802Y^ also exhibited elevated basal kinase activity (p-T899) but this was always weaker than GCN2^M802F^. Structural modelling of the GCN2^M802F^ mutant (Fig 8F) suggested that the bulkier Phe802 causes a substantial clash with the Tyr-down position of Tyr651, indirectly stabilising the active Tyr-up position through direct interactions with Leu640 in the RS3 space and thereby promoting formation of an active back-to-back dimer.

**Figure 9.**
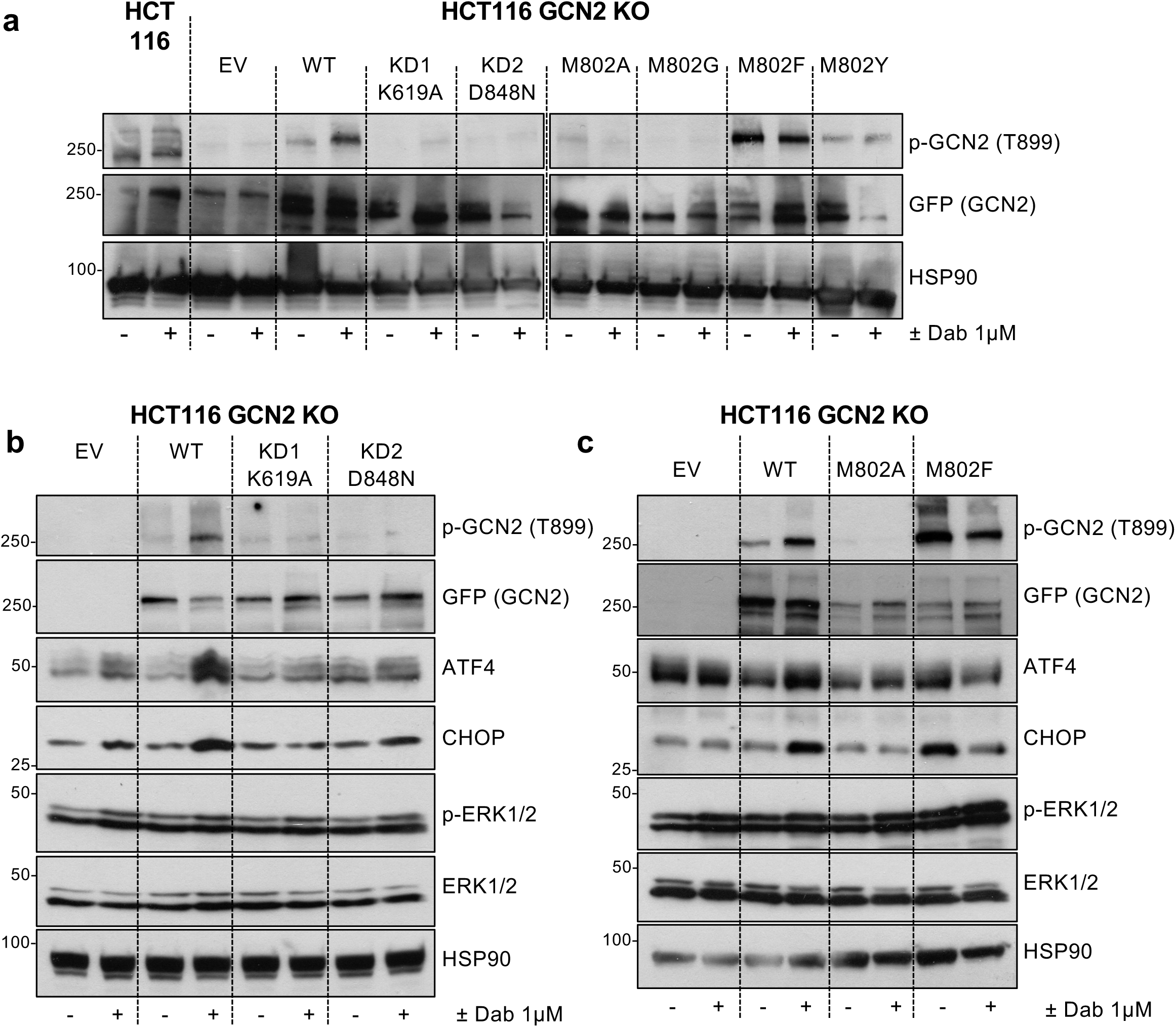
Kinase dead and Gatekeeper mutations abolish Dabrafenib-induced activation of GCN2 and the ISR. **a.** HCT116 GCN2 knockout (KO) cells (clone g1 6) were transiently transfected with either empty vector (EV, EGFP-C3) or constructs expressing wildtype WT EGFP-GCN2, two kinase dead mutants (KD1, K619A or KD2, D848N) or one of four different gatekeeper mutations (M802A, M802G, M802F or M802Y). After 24 hours, cells were treated with either DMSO (-) or 1µM Dabrafenib (+) for 4 hours. HCT116 wild type cells were also treated with drug. Whole cell lysates were separated by SDS-PAGE and immunoblotted with the indicated antibodies. **b.** HCT116 GCN2 KO cells were transfected with either empty vector (EV), wildtype (WT) or the KD1 or KD2 kinase dead mutants; treatments and western blots were performed as above. **c.** HCT116 GCN2 KO cells were transfected with either empty vector (EV), wildtype (WT) or one of the gatekeeper mutants, M802A and M802F; treatments and western blots were performed as above. All experiments are representative of at least 3 replicates.

Supporting these findings, GCN2 KO HCT116 cells exhibited a very weak induction of ATF4 in response to Dabrafenib (Fig 5C & 5D and Fig 9B); this was rescued by expression of WT EGFP-GCN2, but not by the kinase-dead mutants EGFP-GCN2^K619A^ or EGFP-GCN2^D848N^. Expression of EGFP-GCN2^M802A^ also failed to restore Dabrafenib-induced ATF4 and CHOP expression whereas the constitutively active EGFP-GCN2^M802F^ mutant was sufficient to increase expression of ATF4 and CHOP even in the absence of Dabrafenib; indeed, Dabrafenib actually reduced the activity of the EGFP-GCN2^M802F^ mutant. Together these results allow us to conclude that RAFi bind directly to the canonical GCN2 kinase domain to drive paradoxical GCN2 activation, which is required for RAFi to activate the ISR in cells.

During the course of our studies we observed that GCN2iB, an ATP-competitive GCN2 inhibitor ^50^ increased ATF4 expression in a GCN2-dependent fashion (Fig 5C). We compared the effects of GCN2iB with A-92 that we had previously used as a GCN2 inhibitor (Fig 4C, Supplementary Fig 4A). GCN2iB alone caused a striking concentration-dependent increase in ATF4 expression that peaked at 10-30nM and declined at 100-300nM. This response mirrored the increase in pT899-GCN2 and was almost completely lost in GCN2 KO cells indicating that GCN2iB was driving paradoxical activation of GCN2 (Fig 10A). In contrast, whilst A-92 increased p-T899-GCN2, it failed to increase ATF4 expression. To understand the contrasting effects of these two GCN2 inhibitors, we again generated structural models. In common with Dabrafenib, GCN2iB was predicted to bind to GCN2; indeed, there are remarkable similarities in the binding mode of Dabrafenib (Fig 8C) and GCN2iB (Fig 10B); both occupy the RS3 space, stabilise Tyr651 in a Tyr-up position and assemble the R-spine to promote an active back-to-back dimer. In contrast, A-92 is a Type I inhibitor (like GDC-0879); in two different binding ‘poses’ it failed to reach into the RS3 space so the R-spine is not assembled and GCN2 is not activated (Fig 10C & 10D). Thus, GCN2iB, like Dabrafenib, can act as a GCN2 activator at low doses before inhibiting GCN2 dimers at high doses, whereas A-92 acts predominantly as a GCN2 inhibitor. This additional insight explains why A-92 is effective as a GCN2 inhibitor and prevented Dabrafenib-induced ISR activation (Fig 4C).

**Figure 10.**
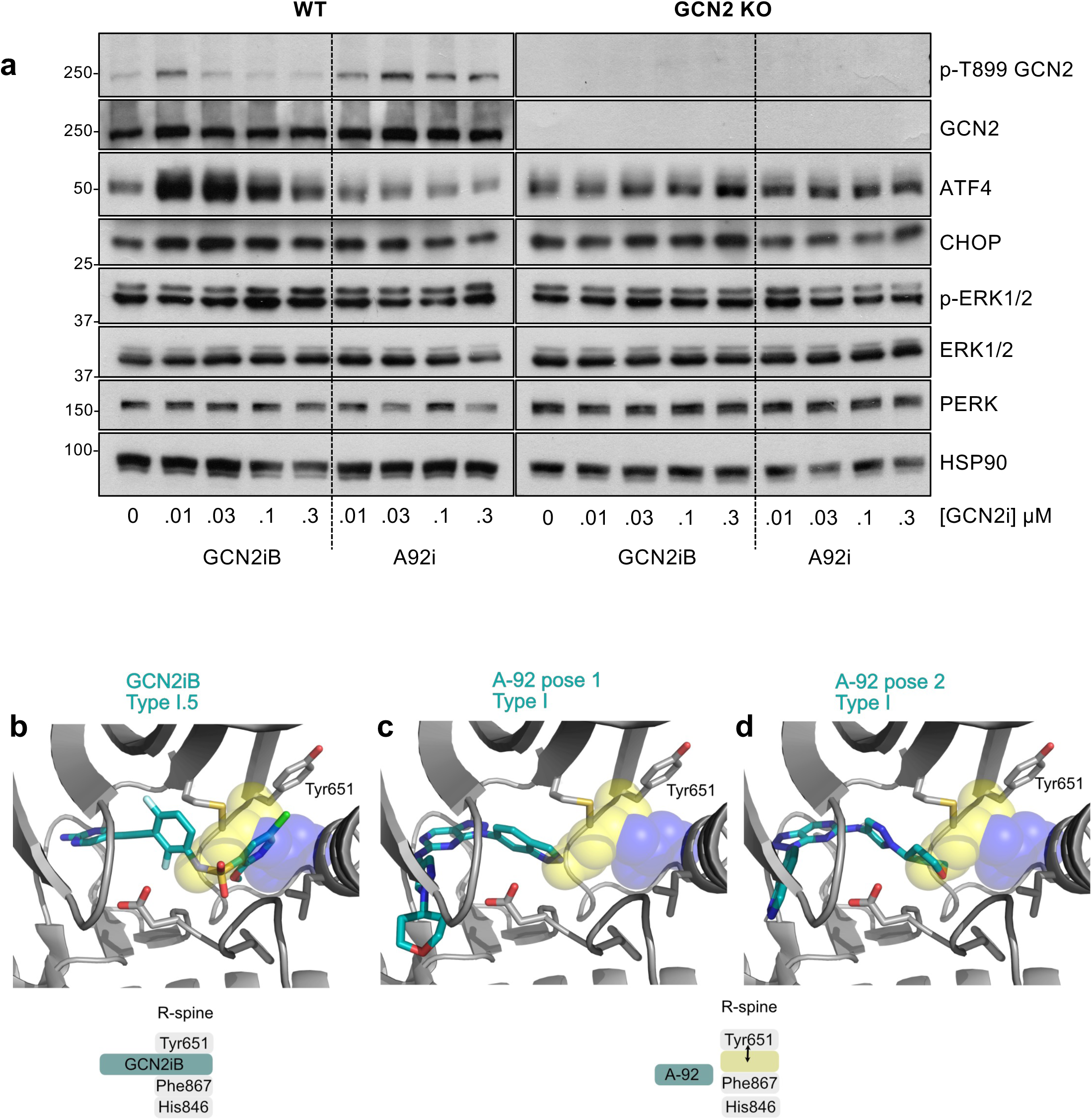
The GCN2 inhibitor GCN2iB drives paradoxical activation of GCN2 whereas A-92 does not. **a.** Wildtype (WT) or GCN2 knockout (KO) HCT116 cells were treated with the indicated concentrations of either GCN2iB or A-92 for 4 hours. Whole cell lysates were separated by SDS-PAGE and immunoblots were probed with the indicated antibodies. Blots from a single experiment representative of 3 independent experiments are shown. **b-d.** Structural models of GCN2iB and A-92 inhibitors. The Tyr-down position and RS3 space are shown as yellow and blue space filling spheres, respectively. **b**. Crystal structure of GCN2 in complex with GCN2iB (PDB code 6N3O). **c and d.** Models of GCN2 in complex with A-92, generated using molecular docking against PDB code 6N3O. Two alternative Type I binding poses are shown.

## Discussion

Whilst studying paradoxical activation of RAF in cancer cells, we found that the majority of RAF inhibitors simultaneously drive activation of GCN2 in cells, resulting in phosphorylation of eIF2α and activation of the ISR. We discovered this due to the unexpected ability of RAFi to drive a rapid inhibition of DNA replication accompanied by loss of Claspin and CDC25A (Fig 2B& 2C) two key regulators of the DNA damage response^51^. However, RAFi did not activate markers of DNA damage such as pS139-H2AX or p-S345 CHK1, the kinase that normally promotes degradation of CDC25 following DNA damage. MEK1/2 inhibition prevented ERK1/2 activation but did not reverse the antiproliferative effect of RAFi, showing it was independent of MEK1/2-ERK1/2 signalling. Rather, RNA-seq revealed that RAFi treatment led to the ERK1/2-independent activation of the Unfolded Protein Response. Ultimately, comparisons with ER stress agonists and use of PERK inhibitors or PERK KO cells ruled out the UPR *per se* and allowed us to focus on the ISR. Knockdown, KO or inhibition (by A-92) of GCN2 and antagonism of p-eIF2α (by ISRIB) prevented the RAFi-induced expression of ATF4, CHOP and their target TRIB3; notably, A-92 and ISRIB also reversed the loss of CDC25A and the inhibition of DNA replication indicating that the antiproliferative effects of RAFi likely reflect activation of the ISR and attenuation of translation.

To investigate the underlying mechanisms we focused on Dabrafenib, Encorafenib, LY-3009120 and GDC-0879. In cells GCN2 activation/ATF4 expression was apparent at 100nM Dabrafenib, Encorafenib or LY-3009120, peaked at 0.5-1µM, with inhibition at 10µM, exhibiting a bell-shaped or Gaussian concentration-response curve. The same response was seen for activation of full-length, purified dimers of recombinant human GCN2 *in vitro* indicating that these inhibitors were binding directly to GCN2 to activate it. In contrast, GDC-0879 failed to activate GCN2 in cells or *in vitro* and served as a useful control. Gatekeeper mutations (GCN2^M802A/G^) in the ATP-binding pocket rendered GCN2 insensitive to Dabrafenib in cells, confirming that RAFi bound to the canonical kinase domain of GCN2 and not the pseudokinase or HisRS domains, which harbour more cryptic ATP-binding sites. Whilst expression of WT GCN2 reconstituted RAFi-induced ATF4 and CHOP expression in GCN2 KO cells, the GCN2^M802A^ mutant and two different kinase-dead mutants (GCN2^K619A^ or GCN2^D848N^) failed to do so, indicating that binding to the ATP-binding pocket, and activation of the GCN2 kinase domain is required for RAFi to activate the ISR.

These results are highly reminiscent of the well-known paradoxical activation of wild type RAF dimers by RAFi ^52^. In common with WT RAF proteins GCN2 functions as back-to-back dimers ^38,39^. Our modelling suggests that GCN2 cross-dimer interactions between the side chains of Tyr651 and Tyr652 are central in the dimeric interface with RAFi-induced kinase activation dependent on Tyr651 adopting the ‘Tyr-up’ position which occupies RS4 to assemble the R-spine and promote back-to-back dimerization and activation^43^; such allosteric activation via the αC/μ4 region is very common in kinases^53, 54^. Our structural models of GCN2-RAFi complexes revealed that both Dabrafenib and LY-3009120 bind to GCN2 in a manner that assembles the R-spine with Tyr651 in the up position; in contrast, GDC-0879 did not, consistent with its failure to activate GCN2 and the ISR in cells. We also observed that low doses of GCN2iB drove paradoxical activation of GCN2 and the ISR whereas another GCN2 inhibitor, A-92, did not. Our structural models suggest that GCN2iB binds to GCN2 in a very similar manner to Dabrafenib to assemble the R-spine whereas A-92 does not. This likely explains why A-92 acted as a GCN2 inhibitor to prevent RAFi-induced activation of the ISR in our studies (Fig 4C). These results confirm and extend a previous report that low doses of GCN2iB could activate GCN2 and this could be inhibited by A-92 ^55^. Based on these results A-92 might seem to be the preferred choice for inhibition of GCN2 in cell-based studies but this is made more complex because A-92 has also been shown to activate PERK in cells ^56^. However, A-92-driven PERK activation was only observed at very high doses of A-92 (10-40µM), far in excess of the doses used herein to inhibit GCN2. Indeed, the high degree of homology between the EIFA2Ks is such that PERK inhibitors can also activate GCN2 in cells and *in vitro*, as can the PKR inhibitor C16 ^56^.

Based on our results and the precedent of RAFi driving paradoxical activation of RAF dimers we propose that at low concentrations RAFi and GCN2iB bind to one protomer of GCN2 and inhibit it, but in doing so they drive allosteric activation of the drug-free, ATP-bound dimer partner through the interplay between Tyr651 and Tyr652 at the dimeric interface. This results in activation of GCN2 in cells, manifest as eIF2α phosphorylation and ATF4 expression. Attempts to formally test this model using mutations that should disrupt GCN2 dimerization are hampered by the fact that such mutations destabilise expression of full length GCN2. Nonetheless, the bell-shaped concentration response for GCN2 activation in cells and *in vitro*, the mutagenesis of the GCN2 kinase domain and expression in GCN2 KO cells and our structural modelling are all entirely consistent with this hypothesis. Notably, whilst both GCN2 and RAF act as dimers, GCN2 is much larger and more complex than the RAF proteins and contains several other domains involved in inhibitory interactions that must be broken for activation ^41^. Whether activation of GCN2 through the back-to-back interface is sufficient for full activation, as is seen with RAF dimers and RAFi, remains to be seen. In a separate study our collaborators have employed HDX-MS to analyse GCN2 activation by RAFi and GCN2iB^57^. As well as changes at the ATP binding site, consistent with our modelling, they have observed propagation of allosteric changes beyond the kinase domain; this perhaps suggests that RAFi or GCN2iB binding in the kinase domain may promote a pseudo-active conformation that initiates further allosteric changes to drive full activation.

In *in vitro* kinase assays we observed a weak increase in eIF2α phosphorylation when it was added to GCN2 kinase reactions as a substrate; however, this was very modest when compared to the strong activation induced by excess ATP. This may be because RAFi bind to the GCN2 kinase domain (supported by modelling, confirmed with ‘gatekeeper’ mutations) and so compete with substrate (eIF2α) binding. Alternatively, RAFi-induced GCN2 autophosphorylation may not feature the same molecular mechanism as substrate phosphorylation and further changes or accessory factors may be required for efficient eIF2α binding or phosphorylation.

Prior studies reported that Dabrafenib^58^ and Vemurafenib^59^ could drive ATF4 expression but the underlying mechanism was not defined. In this study, we report activation of GCN2-ISR with 8 out of 10 chemically-distinct RAFi, suggesting that this is a property common to RAFi as a group, rather than an isolated ‘off-target’ effect of a single RAFi. Both the clinically approved Group 1 RAFi (Vemurafenib, Dabrafenib and Encorafenib) and Group 2 and paradox breaker RAFi activated the ISR to the same extent as ER stress (PERK activators) or histidine starvation (GCN2 activator); only GDC-0879 and AZ628 failed to do so. Furthermore, we observed RAFi-driven activation of GCN2-ISR in tumour cells with wild type BRAF, mutant BRAF, mutant KRAS and immortalised MEFs with no ERK pathway mutation. Moreover, activation of GCN2 is not confined to EIF2AK inhibitors or RAFi; in the course of our work certain EGFR tyrosine kinase inhibitors including Erlotinib and Neratinib have also been shown to activate GCN2 in glioblastoma cell lines ^60^. However, in this case the doses of EGFRi required for GCN2 activation were substantially in excess of that required to inhibit EGFR. This contrasts with the effects reported here, where the majority of RAFi tested, including Dabrafenib, Encorafenib and Vemurafenib, activated GCN2 and the ISR over the same concentration range at which they activated RAF and the ERK1/2 pathway. This raises important questions about the extent to which ISR activation influences the clinical response to BRAF inhibitors. In cancers harbouring BRAF^V600E/K^ ISR activation will be coincident with ERK1/2 inhibition, itself a pro-survival pathway^61^, and ISR activation may be a route to RAFi resistance. In cells lacking BRAF mutations coincident ERK1/2 and GCN2-ISR activation may contribute to adventitious benign tumour growth. More generally, it suggests that non-kinase pocket inhibitors of RAF such as dimerization inhibitors might be more desirable to avoid paradoxical activation of GCN2 and the ISR as well as RAF and the ERK1/2 pathway.

## Materials and Methods

### Reagents and Cell Lines

The source of all reagents and cell lines utilised are detailed in Table S1. Cells were grown in DMEM (HCT116) RPMI1640 (NCI-H358 and HT29) media supplemented with 10% (v/v) fetal bovine serum, penicillin (100U/mL), streptomycin (100mg/mL) and 2mM glutamine. Cells were incubated in a humidified incubator at 37°C and 5% (v/v) CO2. All cell lines were authenticated by Short Tandem Repeat (STR) profiling and confirmed negative for mycoplasma prior to experiments commencing. All reagents were from Gibco, Thermo Fisher Scientific (Paisley, UK).

### High content microscopy ^26,27^

Cells were cultured in CellCarrier-96 plates (PerkinElmer). After the drug treatment, cells were harvested, fixed with 4% (v/v) formaldehyde/PBS and permeabilised with 0.2% Triton X-100 for 10 minutes. Cells were blocked for 1 hour with 2.5% goat serum (v/v) in 2% BSA/PBS (v/v) at room temperature. Cells were then incubated with primary antibody diluted in 2% BSA/PBS overnight at 4°C. Background control wells were treated with 2% BSA/PBS, without the addition of primary antibody. Cells were washed three times with PBS and then incubated with Alexa Fluor secondary antibodies (1:500) and DAPI (1 ug/ml) in 2% BSA/PBS for 1 hour. Cells were washed with PBS before imaging using an INCELL Analyzer 6000 Microscope (GE Healthcare), imaging six fields per well. Image analysis to determine the mean signal intensity was performed using INCELL Analyzer software. For Edu analysis, cells were treated as described in the figure legends and 1 hour prior to harvest incubated with 10μM 5-ethynyl-2-deoxyuridine (EdU; Click-iT EdU HCS Kit, ThermoFisher, Loughborough, UK). Fixed and processed as above and EdU detected according to the manufacturer’s protocol.

### EdU incorporation by flow cytometry

Cells were treated as described in the figure legends and 1 hour prior to harvest incubated with 10μM 5-ethynyl-2-deoxyuridine (EdU; Click-iT EdU Flow Cytometry Kit, ThermoFisher, Loughborough, UK). Cells were harvested by trypsinisation and fixed with 4% paraformaldehyde/PBS for 10 min at room temperature. EdU was detected following the manufacturer’s instructions, and cells were resuspended in 1µg/mL DAPI/PBS (Sigma-Aldrich, Dorset, UK). DAPI and EdU staining was assessed with a FACS LSRII (BD Biosciences, Oxford, UK), counting 10000 cells per sample. Data was analyzed using FlowJo software (FlowJo, Oregon, USA).

### RNA-seq

At the Babraham Institute NCI-H358 cells were treated DMSO, 1µM Dabrafenib (Dab), 30nM Selumetinib (Sel) or 1µM Dabrafenib + 30nM Selumetinib (Dab+Sel) for 4hrs or 24 hrs. Four independent biological repeats were prepared making a total of 32 samples. Cells were lysed in RLT buffer, with the addition of β-mercaptoethanol. Lysates was disrupted using Qiashredder columns (Qiagen) and RNA prepared as described in the manufacturers protocol (Qiagen, RNeasy plus) with samples eluted in RNase-free water. Samples were analysed by Nanodrop spectrophotometer before being shipped to Plexxikon Inc., CA. Post shipping, Nanodrop analysis was repeated and fragment analysis was performed for quality control purposes at Plexxikon before shipping to the UC Berkeley QB3 Genomics Facility. Strand specific RNA-seq libraries were prepared from poly-A enriched RNA using a Kapa Biosystems Kit and then sequenced (150 bp paired end reads) on a NovaSeq S4 flow cell (QB3 Genomics, UC Berkeley, Berkeley, CA, RRID:SCR_022170). Reads were trimmed for quality and to remove adapter sequences using fastp version 0.23.4^62^. Trimmed reads were then quantified with salmon (v1.10.1) using the hg38 transcriptome index with full decoy sequences downloaded from RefGenie^63,64^. The R package tximeta was used to calculate offsets, annotate and summarize expression to the gene level^65^. Then, the edgeR package was used to filter out genes not expressed in any sample, estimate dispersion, fit a generalized linear model and identify differentially expressed genes^66^. Finally, the camera function from the limma package was used to test whether sets of genes (from the molecular signatures database Hallmark genesets) were highly ranked relative to other genes in terms of differential expression^67^. Data are available at Gene Expression Omnibus (https://www.ncbi.nlm.nih.gov/geo/) under accession number GSE271504

### Preparation of cell lysates for SDS-PAGE and Western blotting

Culture medium from cells growing on dishes was either removed. Cells were washed with PBS and then and harvested in lysis buffer (20 mM Tris [pH 7.5], 137 mM NaCl, 1 mM EGTA, 1% (v/v) Triton X-100, 10% (v/v) glycerol, 1.5 mM MgCl2, 1 mM Na3VO4, 1 mM PMSF, 20 mM leupeptin, 10 mg/ml aprotinin and 50 mM NaF). Cell extracts were snap frozen, thawed and cleared by centrifugation. Supernatant protein concentration was determined by Bradford protein assay (Bio-Rad) and absorbance measured at 562 nm using a PHERAstar FS plate reader (BMG Labtech, Aylesbury, UK). Samples were prepared for SDS-PAGE by boiling for 5 min in 1 x Laemmli sample buffer (50 mM Tris-HCl (pH 6.8), 2% (w/v) SDS, 10% (v/v) glycerol, 1% (v/v) β-mercaptoethanol, 0.01% (w/v) bromophenol blue).

### SDS-PAGE and Western blotting

Lysates were separated by SDS-PAGE (Mighty small II gel apparatus, Hoefer, Massachusetts, USA). Polyacrylamide gels consisted of a resolving phase of 8-16% (w/v) acrylamide (37.5:1 acrylamide:bisacrylamide, 2.7% crosslinker; Bio-Rad, Watford, UK), 0.375 M Tris-HCl (pH 8.8), 0.2% (w/v) SDS (Bio-Rad, Watford, UK), 0.1% (w/v) ammonium persulfate, 0.1% TEMED (Bio-Rad, Watford, UK) and a stacking phase of 4.5% (w/v) acrylamide (37.5:1 acrylamide:bisacrylamide, 2.7% crosslinker), 0.125 M Tris-HCl (pH 6.8), 0.2% (w/v) SDS (Bio-Rad, Watford, UK), 0.1% ammonium persulfate, 0.125% TEMED. Gels were run using running buffer (0.2 M glycine, 25 mM Tris, 0.1% (w/v) SDS) and a current of 15 mA per gel for 3-4 hours. Gels were then blotted by wet transfer (Bio-Rad, Watford, UK) to methanol activated PVDF (Immobilon-P Membrane, Merck Millipore, Watford, UK) using transfer buffer (0.2 M glycine, 25 mM Tris, 20% (v/v) methanol) and a current of 300 mA for 90-180 min. Membranes were blocked in 5% milk/TBST (5% (w/v) non-fat powdered milk, 10 mM Tris-HCl (pH 8.0), 150 mM NaCl, 0.1% (v/v) Tween-20) for 1 hour at room temperature. Membranes were incubated with primary antibodies as recommended in 5% milk/TBST or 5% BSA/TBST overnight at 4°C with agitation. Antibodies are detailed in Table S1. Membranes were then washed in TBST for 3 x 10 min, and incubated with horseradish peroxidase-conjugated secondary antibodies (Bio-Rad, Watford, UK) diluted 1:3000 in 5% milk/TBST, or fluorescently labelled secondary antibodies diluted 1:15000 (Cell Signaling Technology, NEB, Hitchin, UK), for 1 hour at room temperature. Membranes were again washed for 3 x 10 min in TBST. Detection was performed using Amersham ECL Western Blotting Detection Reagent (Cytiva), X-ray film and Compact X4 film developer (Xograph, Gloucestershire, UK). Quantification of fluorescently labelled membranes was performed using the Odyssey Infrared imaging system (LI-COR, Cambridge, UK).

### GCN2 knockout by CRISPR/Cas9-mediated gene editing

Plasmids containing guide RNAs (gRNAs) to *GCN2;* hsEIF2AK4_guide1_pSpCas9(BB)-2A-Puro (MP1) and hsEIF2AK4_guide2_pSpCas9(BB)-2A-Puro (MP1) were a kind gift from Dr H Harding, Cambridge University and were subcloned into a pSpCas9(BB)-2A-GFP genome editing vector, which was a gift from Feng Zhang (Addgene plasmid #48138). Cells were transfected with the *GCN2* gRNA containing Cas9 plasmids using jetPrime (Polyplus Transfection, Illkirch, France). Transfection was monitored by GFP expression and single GFP positive/DAPI negative cells were sorted in to 96 well plates using a 100 µm nozzle on a BD FACSARIA III cell sorter (BD Biosciences, Oxford, UK). Clones of interest were identified by Western blot screening for absence of GCN2.

### DNA sequencing to confirm absence or presence of GCN2 mutations

Genomic DNA was extracted from HCT116 (control untransfected) and all WT and KO clones using Qiagen quick extract according to the manufacturer’s protocol. Genomic DNA flanking the CRISPR guide binding site was amplified by PCR using OneTaq (NEB, Hitchin, UK) according to the manufacturer’s instructions and using the primers indicated in table S1. The products generated were then cloned into the TOPO-TA cloning vector (Thermo Fisher Scientific, Loughborough, UK) following the manufacturer’s instructions. The resulting constructs were used to transform chemically competent DH5α (NEB, Hitchin, UK), 4-5 of the resulting clones derived from DNA for each cell line were sent for sequencing (Genewiz, Bishop’s Stortford, UK).

### GCN2 Kinase Assays

Expression and purification of recombinant human GCN2 and eIF2α was conducted as described previously ^68^. 10 nM GCN2 was incubated at 30 °C for 20 min with 2 µM eIF2α and 10 µM ATP. Samples were then quenched via the addition of SDS-PAGE loading buffer and heating to 95 °C for 3 minutes. Samples were separated on by SDS-PAGE (5-12% Bis-Tris Gel), run at 180 V for 35 min in MES buffer (50 mM MES (2-[N-morpholino]ethanesulfonic acid), 50 mM Tris base, 1 mM EDTA. 0.1% (w/v) SDS). Gels were then briefly incubated in 20% ethanol solution prior to a transfer to a nitrocellulose membrane using the iBlot 2 Transfer system (ThermoFisher Scientific). Membranes were processed as above except LICOR secondary antibodies were used.

### Modelling of GCN2-inhibitor complexes

GCN2-ligand complex models were generated using the Webina implementation of AutoDock Vina ^69^ based on relevant crystal structures (Dabrafenib and GDC0879 used GCN2 bound to a Type I.5 inhibitor, PDB code 6N30 ^37^; LY3002190 used GCN2 bound to a Type II inhibitor, PDB code 6N3L ^37^). Docking poses were selected based on closest match to the equivalent BRAF crystal structures. Met802Phe structure was modelled in PyMOL, based on the AlphaFold2 model of active GCN2, taken from KinCore database ^70,71^. The structure of inactive, yeast GCN2 was used to model the position of Tyr651 in the Tyr-down position ^42^ (PMID: 15964839).

### Transfection

Cells were cultured to approximately 60% confluency. Constructs were then transfected using JetPrime polyplus transfection reagent (Polyplus Transfection, Illkirch, France) or lipofectamine 3000 (ThermoFisher Scientific) according to manufacturer’s instructions. siRNA transfections were performed using RNAiMAX (ThermoFisher Scientific) according to manufacturer’s protocols and using 10nM final siRNA concentration.

### GCN2 expression constructs

GCN2 was amplified from GST-GCN2 (MRC, Dundee PPU-DU67392-proteins-703081) with primers to introduce Xho1 and Xba1 restriction sites at the 5’ and 3’ of the coding regions respectively. The PCR product was cloned into EGFP-C3 (Clontech). Primers for mutagenesis were designed using NEBaseChanger (sequences shown in table S1) and mutagenesis was performed using the Q5 Site-Directed Mutagenesis Kit (New England Biolaboratories). DNA was transformed into stable competent cells (New England Biolaboratories). Whole plasmids were fully sequenced using Plasmidsaurus.

## Acknowledgements

We would like to thank past and present members of the Cook lab for their support and encouragement. We thank Rahul Samant, Ian McGough and Hayley Sharpe (Signalling Programme, the Babraham Institute) and Benedict Cross (PhoreMost) for suggestions and encouragement. We would like to acknowledge the excellent support of the Babraham Institute Science Services including Simon Walker and Hanneke Okkenhaug (Imaging) and Rachael Walker (Flow Cytometry), supported by a Core Capability Grant from the Biotechnology and Biological Sciences Research Council (UKRI-BBSRC). Work in Simon Cook’s laboratory was supported by Plexxikon Inc (AMK and SJC) and Institute Strategic Programme Grants BB/J004456/1, BB/P013384/1 and BB/Y006925/1 from UKRI-BBSRC (SJC and RG). Work in Glenn Masson’s laboratory was supported by a University of Dundee Baxter Fellowship and a TENOVUS Grant T21/01 119615. Work in Patrick Eyer’s laboratory was supported by UKRI-BBSRC grants BB/S018514/1 and BB/X002780/1 while work in Richard Bayliss’s laboratory was supported by Cancer Research UK (C24461/A23302).

## CRediT Author Contribution

Rebecca Gilley: Conceptualisation, Investigation, Formal analysis, Writing - review and editing. Andrew Kidger: Conceptualisation, Investigation, Formal analysis, Writing - review and editing. Graham Neill: Conceptualisation, Investigation, Formal analysis. Paul Severson: Formal analysis. Dominic Byrne: Investigation. Niall Kenneth: Investigation, Writing - review and editing. Gideon Bollag: Resources, Writing - review, Funding acquisition and editing Chao Zhang: Resources, Writing - review, Funding acquisition. Taiana Maia de Oliveira: Conceptualisation, Writing – review and editing. Patrick Eyers: Conceptualisation, Writing - review and editing; Richard Bayliss: Conceptualisation, Writing - review and editing. Glenn Masson: Investigation, Resources, Supervision, Funding acquisition, Writing – review and editing. Simon Cook: Conceptualisation, Resources, Supervision, Funding acquisition, Writing — original draft, review and editing, Project administration.

## Supplementary Information

**Supplementary Figure 1.**
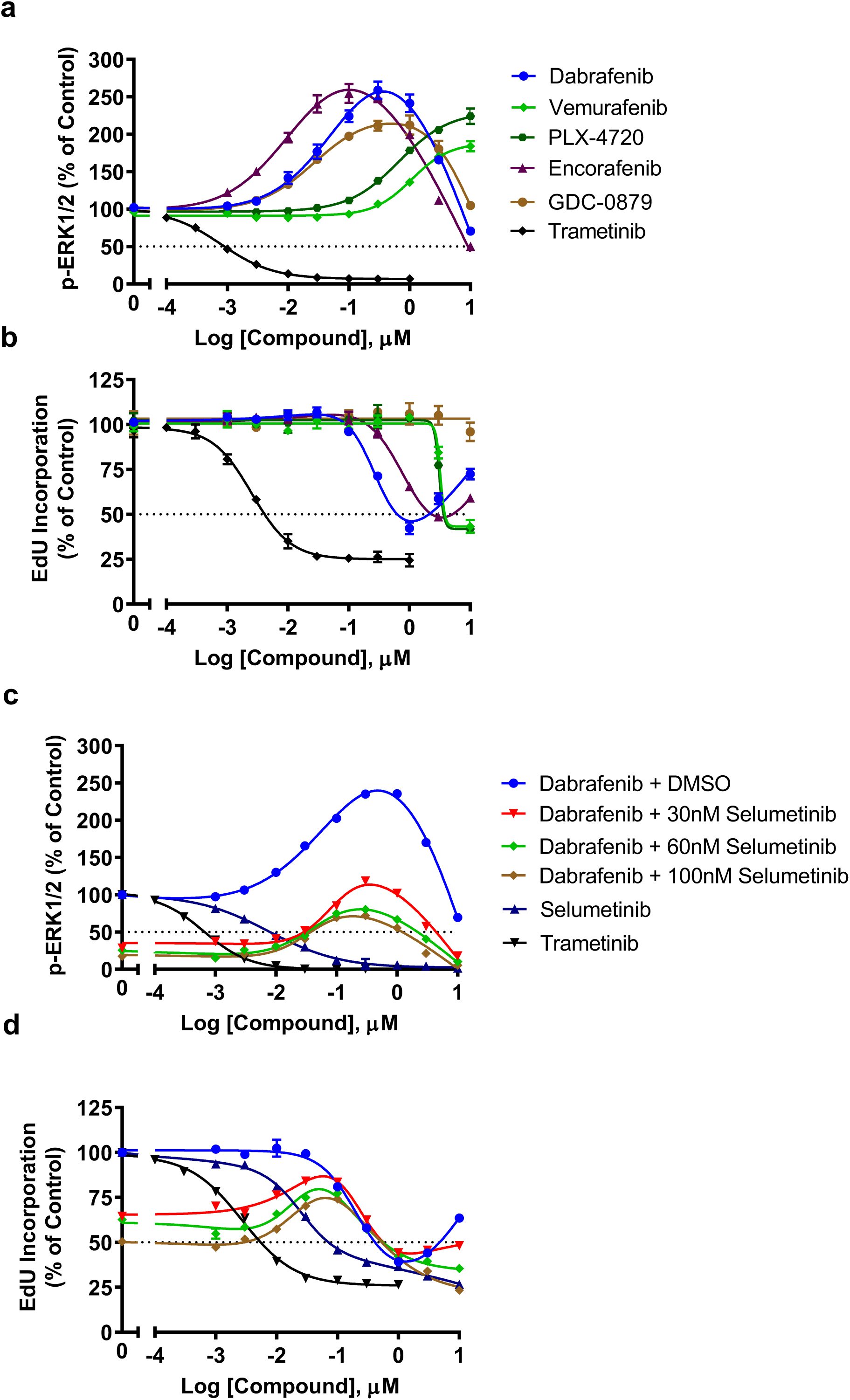
RAF inhibitors inhibit cell proliferation independent of paradoxical ERK1/2 activation. **a & b.** NCI-H358 cells were treated with the indicated concentrations of Dabrafenib, Vemurafenib, PLX-72840, Encorafenib, GDC-0879 or Trametinib for 24 hours with a pulse of 10μM EdU for the final hour. Cells were fixed and permeabilized for EdU detection, immunofluorescence with a p-ERK1/2 antibody and co-stained with DAPI. Mean signal per cell was determined by high-content image analysis of 2000-15000 cells per condition. Normalised mean values ± SEM are shown, n = 3 replicate experiments analysed for p-ERK1/2 (**a**) or intensity of EdU staining (**b**). **c & d.** NCI-H358 cells were treated with 30, 60 or 100nM Selumetinib for 1 hour prior to addition of the indicated concentration of Dabrafenib for 24 hours. Some cells received increasing concentrations of Selumetinib or Trametinib alone for 24 hours. p-ERK1/2 (**c**) and intensity of EdU staining (**d**) were quantified by HCM analysis as above. Results shown are mean ± SEM of n=3 replicate experiments.

**Supplementary Figure 2.**
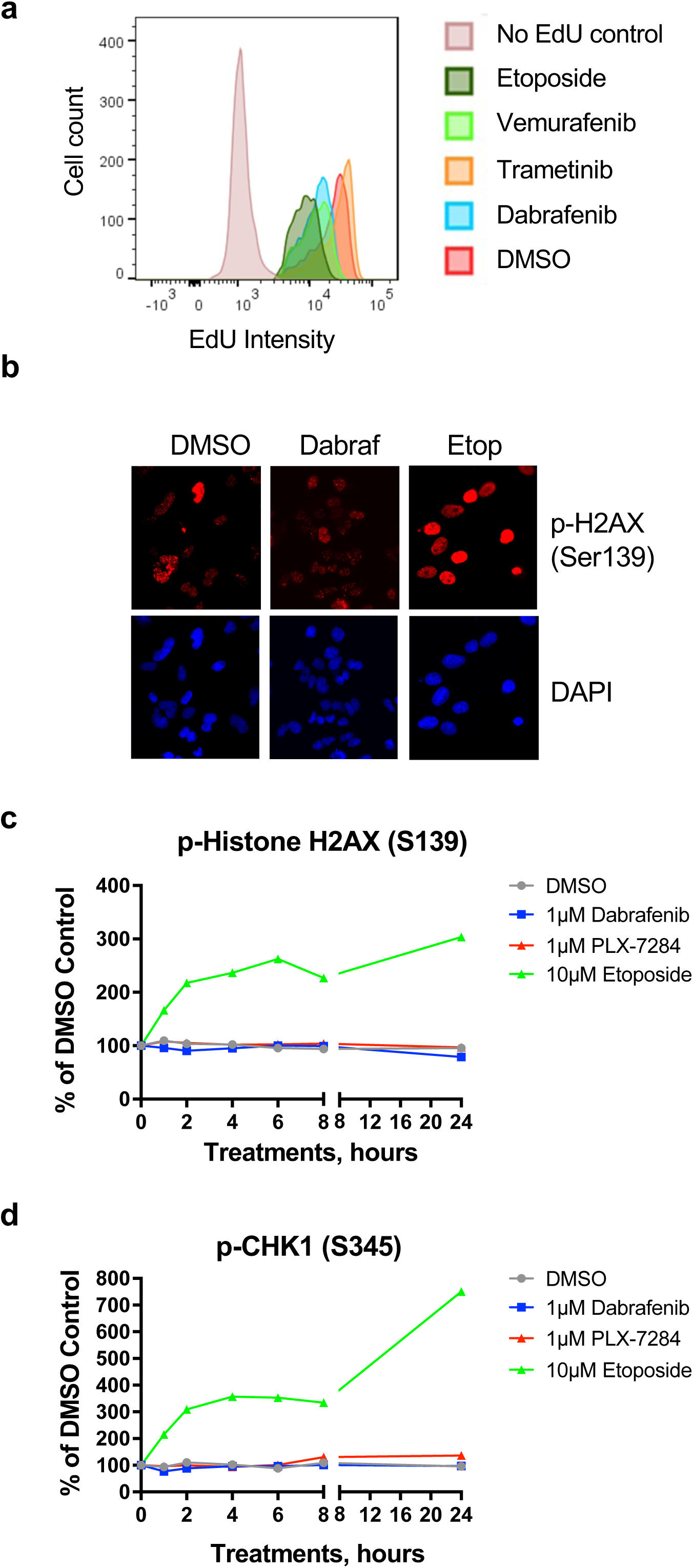
RAF inhibitors drive the rapid inhibition of DNA replication but do not activate the DNA damage response. **a.** NCI-H358 cells were treated with 1µM Dabrafenib, 10µM Vemurafenib, 100nM Trametinib or 10µM Etoposide for 24hrs with a pulse of EdU for the final hour. Cells were harvested and EdU intensity was measured by flow cytometry**. b.** NCI-H358 cells treated with 1µM Dabrafenib or 10µM Etoposide for 24 hours were fixed and permeabilized for immunofluorescence with p-H2AX (Ser139) antibody, co-stained with DAPI and analysed by confocal microscopy. **c & d.** NCI-H358 cells were treated with 1µM Dabrafenib, 1µM PLX-7284 or 10µM Etoposide for up to 24 hours. Cells were then fixed and permeabilized for immunofluorescence with (**c**) p-H2AX(S139) or (**d**) p-CHK1(S345) antibodies. Mean signal per cell was determined by high-content image analysis of 2000-15000 cells per condition. Normalised mean values ± SEM are shown, n = 2.

**Supplementary Figure 3.**
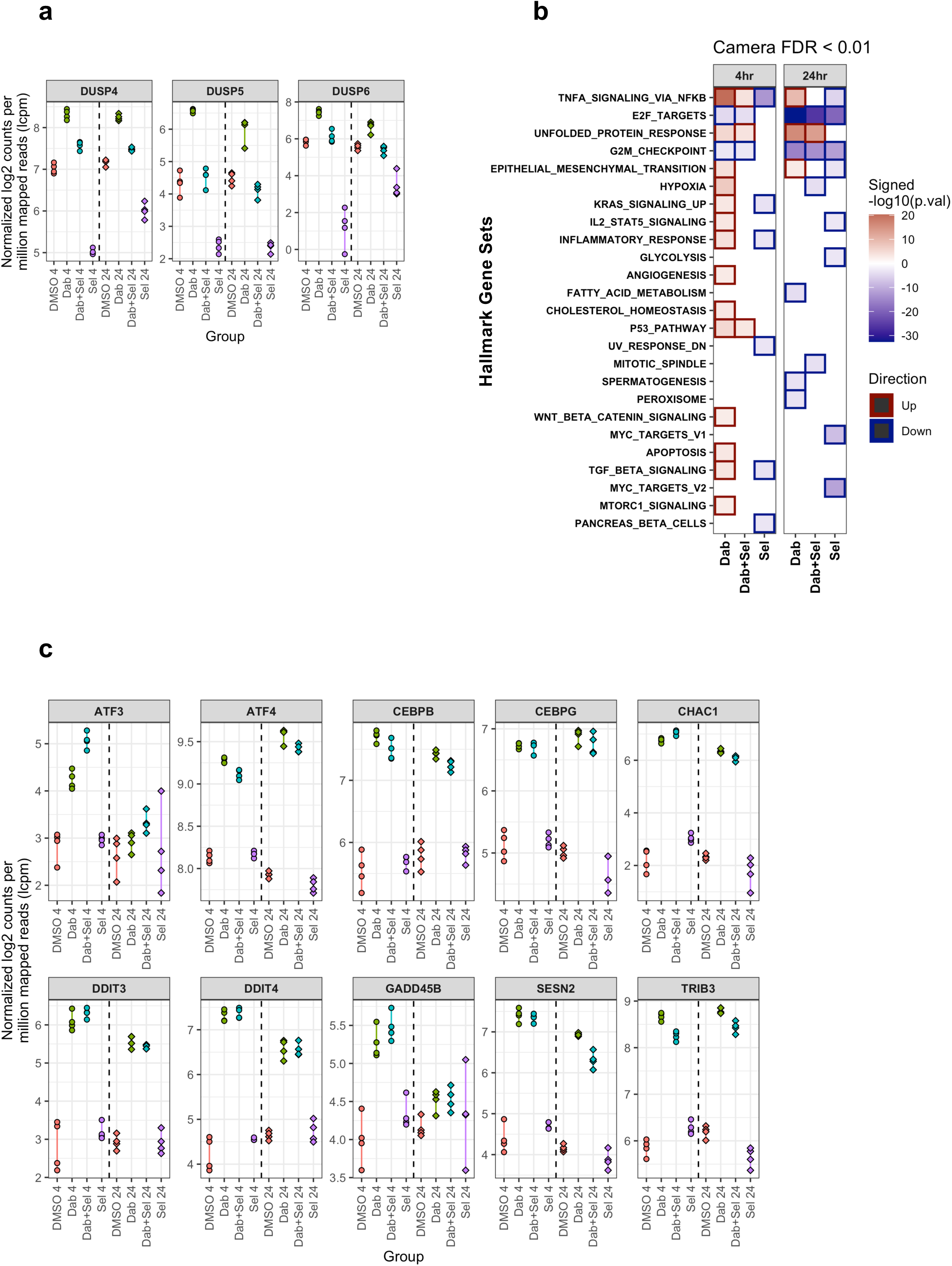
RNA-seq reveals that Dabrafenib activates the unfolded protein response independently of MEK1/2-ERK1/2 signalling. NCI-H358 cells were treated with control (DMSO), 1µM Dabrafenib (Dab), 30nM Selumetinib (Sel) or the combination (Dab+Sel) for 4hrs or 24hrs, with 4 dishes of cells for each condition/timepoint. Samples were processed for RNA-seq as described in the Results and Materials and Methods. **a.** RNA-seq data for the ‘ERK1/2 target genes’ *DUSP4, DUSP5* and *DUSP6* displaying Log2 counts per million mapped reads for each sample, colour-coded by treatment group with symbols representing 4 hours (circles) or 24 hours (diamonds). **b.** RNA-seq data was mapped to the Hallmark genesets. The camera function from the limma package was used to test whether each gene set was highly ranked relative to other genes. The heatmap shows genesets below FDR of 0.01 with tiles colored by significance level (signed, -log10(p.val)). The Unfolded Protein Response geneset was induced by both Dab and Dab+Sel but not Sel alone. **c**. RNA-seq data for ten ‘UPR genes’ displaying Log2 counts per million mapped reads for each sample, colour-coded by treatment group with symbols representing 4 hours (circles) or 24 hours (diamonds). All UPR genes were induced by Dab and Dab+Sel but largely unaffected by Sel alone.

**Supplementary Figure 4.**
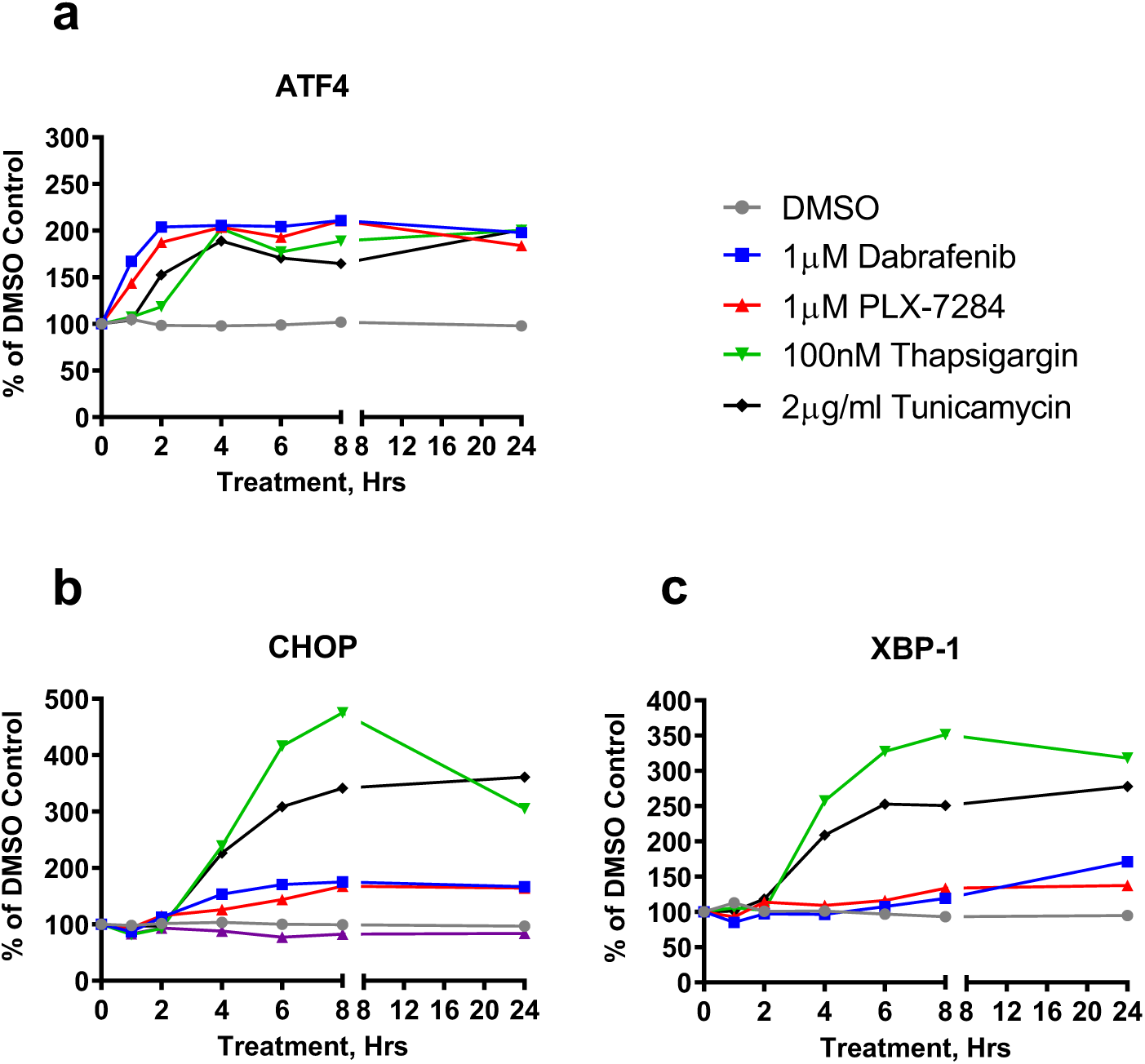
RAFi and ER stressors drive rapid, strong expression of ATF4 but RAFi drive very weak expression of other ER stress markers such as CHOP and XBP-1. NCI-H358 cells were treated with either 1µM Dabrafenib, 1 µM PLX-7284, 100nM Thapsigargin or 2µg/ml Tunicamycin for up to 24 hours. Cells were then fixed and permeabilized for immunofluorescence with antibodies to ATF-4 (**a**) CHOP (**b)** or XBP-1 (**c**). Mean signal per cell was determined by high-content image analysis of 2000-15000 cells per condition.

**Supplementary Figure 5.**
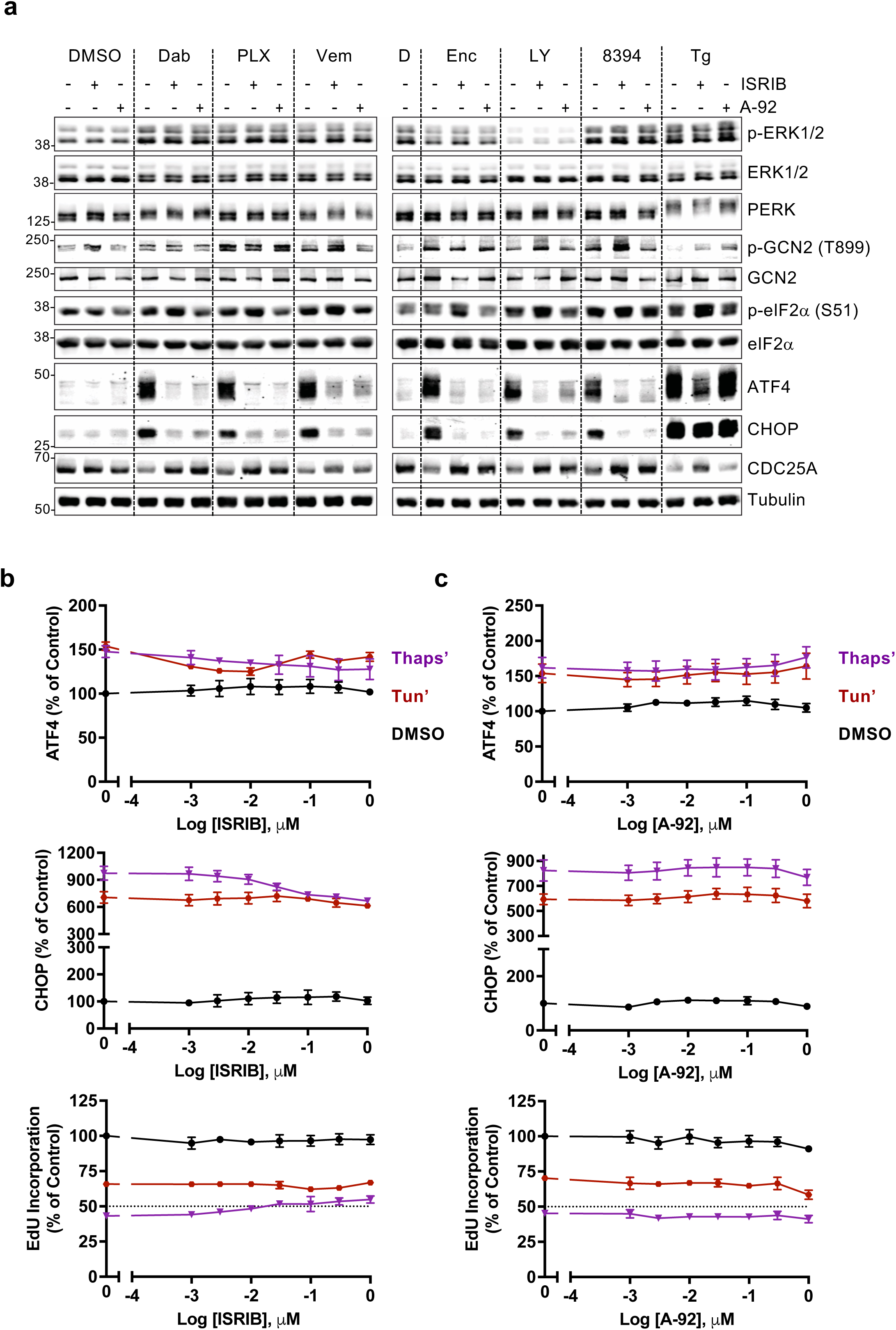
RAF inhibitors, but not ER stressors, require GCN2 kinase activity to activate the ISR. **a.** NCI-H358 cells were treated with DMSO, 300nM ISRIB or 300nM A-92 for 30 mins prior to the addition of either 1μM Dabrafenib, 1μM PLX7284, 10μM Vemurafenib, 3μM Encorafenib, 1μM LY-3009120, 3μM PLX8394 or 100nM Thapsigargin for 4 hours. Whole cell lysates were fractionated and immunoblots were probed with the indicated antibodies and captured by LiCor. Results shown are from a single experiment representative of 3 independent experiments. **b. & c.** NCI-H358 cells were treated with the indicated concentrations of either ISRIB (**b**) or A-92 (**c**) for 30 mins prior to the addition of DMSO, 100nM Thapsigargin or 2µg/ml Tunicamycin. Cells were treated for 7 hours with a pulse of 10μM EdU for the final hour, then fixed and permeabilized for EdU detection, immunofluorescence with ATF4- or CHOP-specific antibodies and co-stained with DAPI. Mean signal per cell was quantified by high-content image analysis of 2000-15000 cells per condition. Normalised mean values ± SEM are shown, from n = 3 replicate experiments.

**Supplementary Figure 6.**
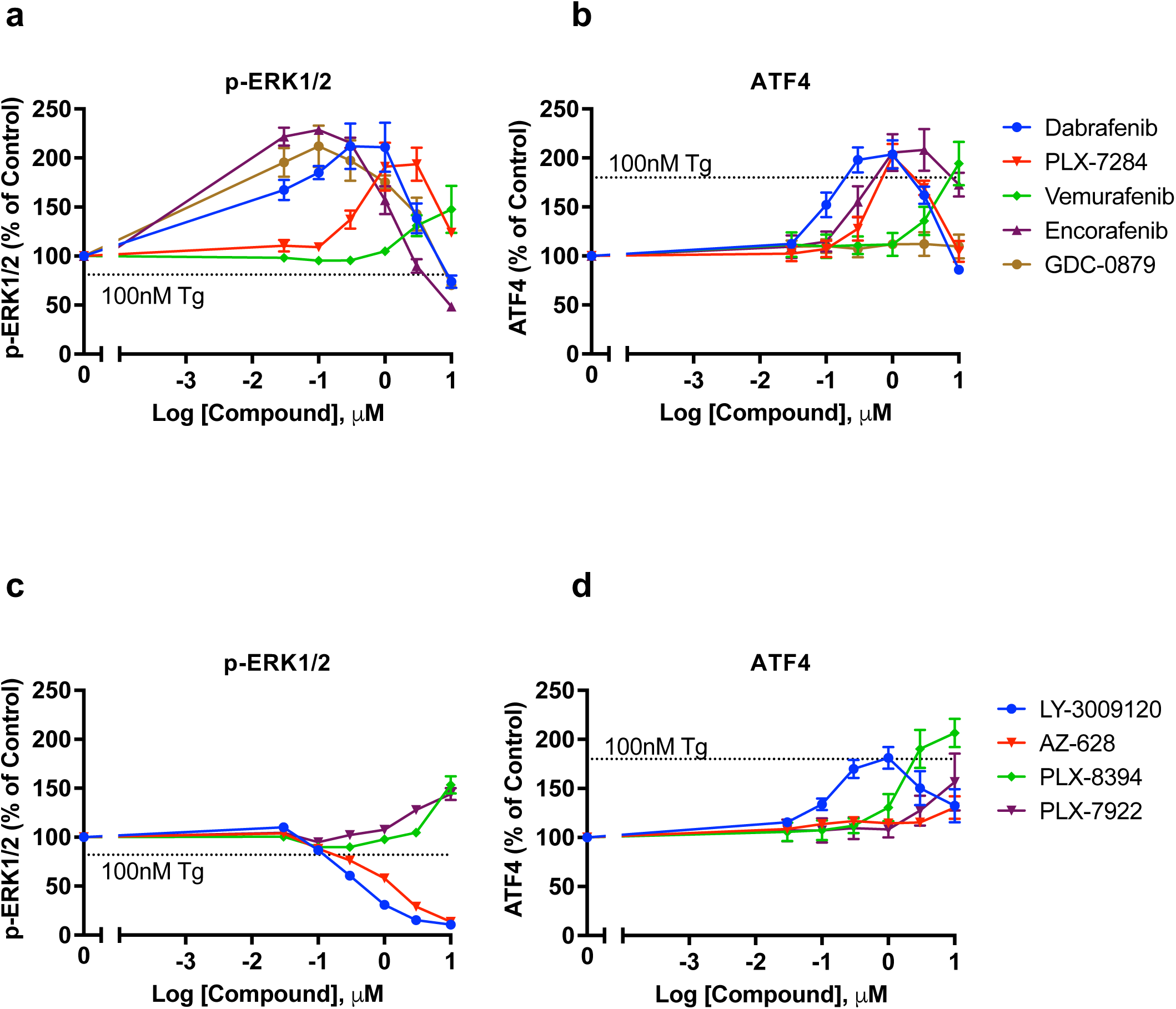
The majority of RAF inhibitors drive paradoxical activation of both ERK1/2 and the ISR. NCI-H358 cells were treated with either (**a & b**) group 1 RAF inhibitors (Dabrafenib, PLX-7284, Vemurafenib, Encorafenib, or GDC-0879) or (**c & d**) group 2 and paradox breaker RAFi (LY-3009120, AZ-628, PLX-8394 or PLX-7922) for 8 hours. Cells were then fixed and permeabilized for immunofluorescence with antibodies to p-ERK1/2 (**a & c**) or ATF-4 (**b & d**). Mean signal per cell was determined by high-content image analysis of 2000-15000 cells per condition. Normalised mean values ± SEM are shown, n = 3

**Supplementary Figure 7.**
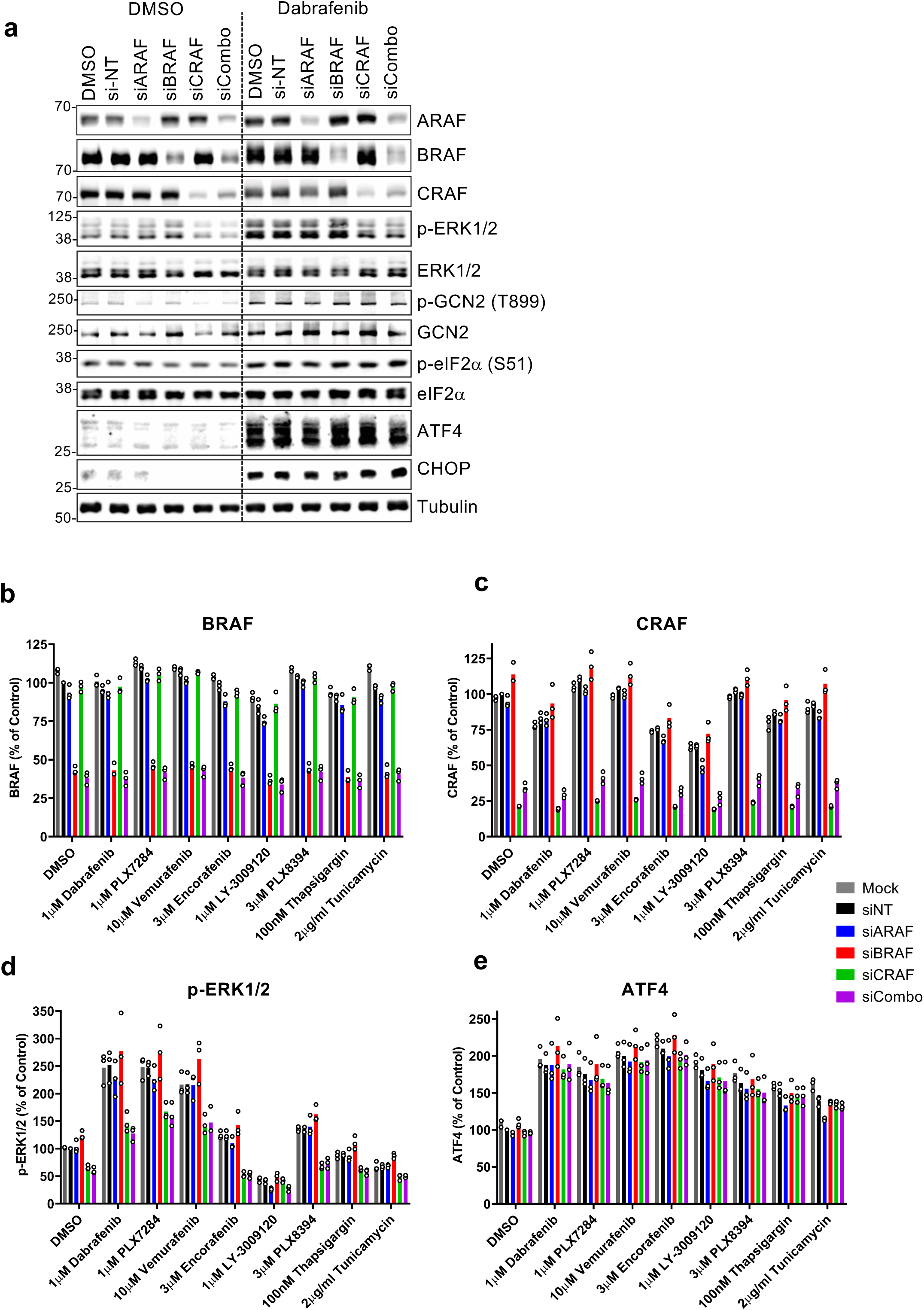
Knockdown of CRAF inhibits Dabrafenib-induced activation of ERK1/2 but not activation of GCN2 and the ISR. **a.** NCI-H358 cells were either mock transfected (MOCK) or transfected with 10nM of siRNA either non targeting (siN-T), targeted to ARAF (siARAF), BRAF (siBRAF), CRAF (siCRAF) or a combination of all (siCombo). After 48 hours cells were treated with either DMSO or 1µM Dabrafenib for 4 hours and whole cell lysates fractionated by SDS-PAGE and immunoblotted with the indicated antibodies. **b-e.** NCI-H358 were transfected as above and after 48 hours were treated with either DMSO, 1μM Dabrafenib, 1μM PLX7284, 10μM Vemurafenib, 3μM Encorafenib, 1μM LY-3009120, 3μM PLX8394, 100nM Thapsigargin or 2μg/ml Tunicamycin for 7 hours. Cells were then fixed and permeabilized for immunofluorescence with antibodies to **b.** BRAF **c.** CRAF **d.** p-ERK1/2 or **e.** ATF4 and co-stained with DAPI. Mean signal per cell was determined by high-content image analysis of 2000-15000 cells per condition. Normalised mean values ± SEM are shown, n = 3 replicate experiments.

**Supplementary Table 1.**
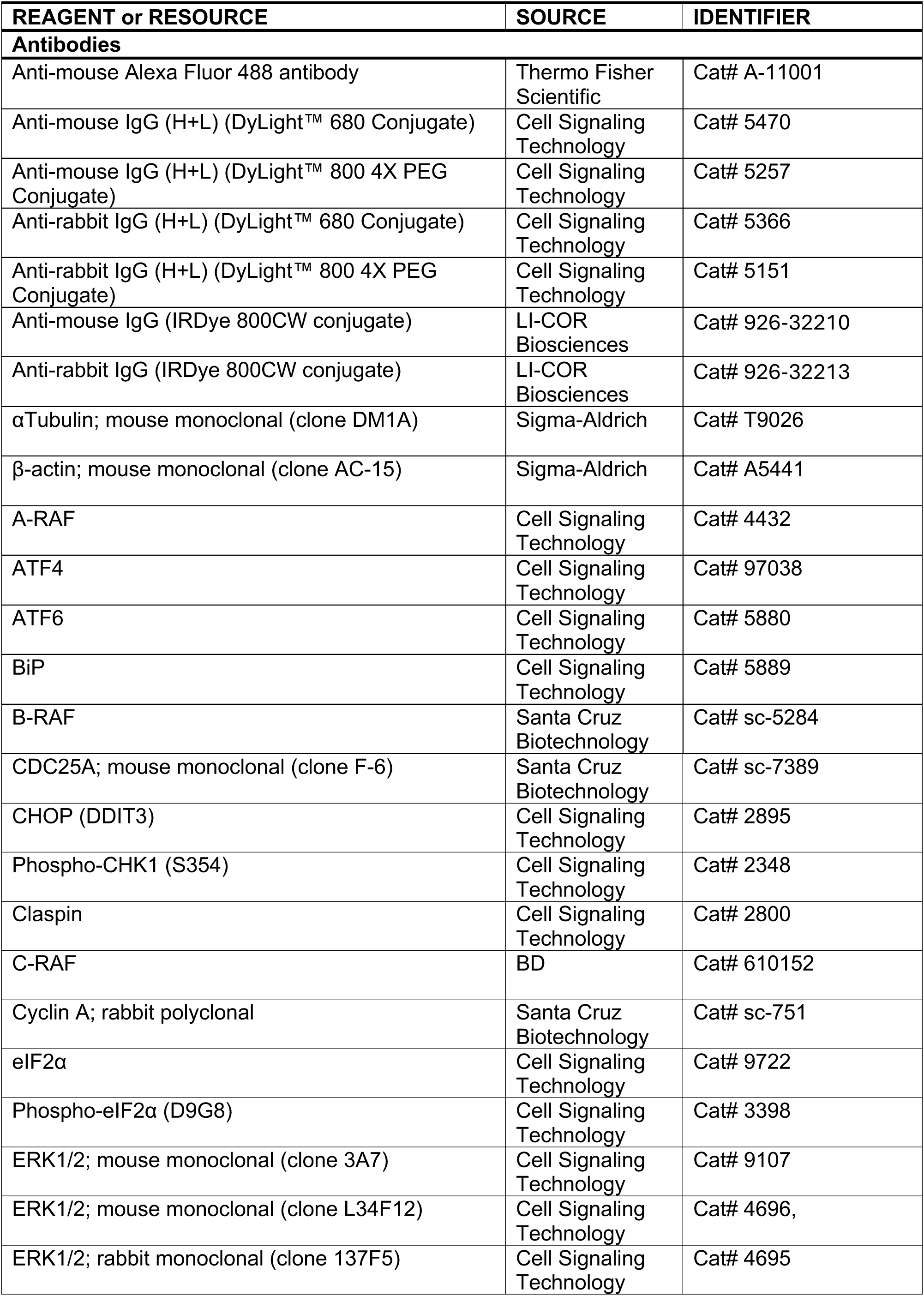
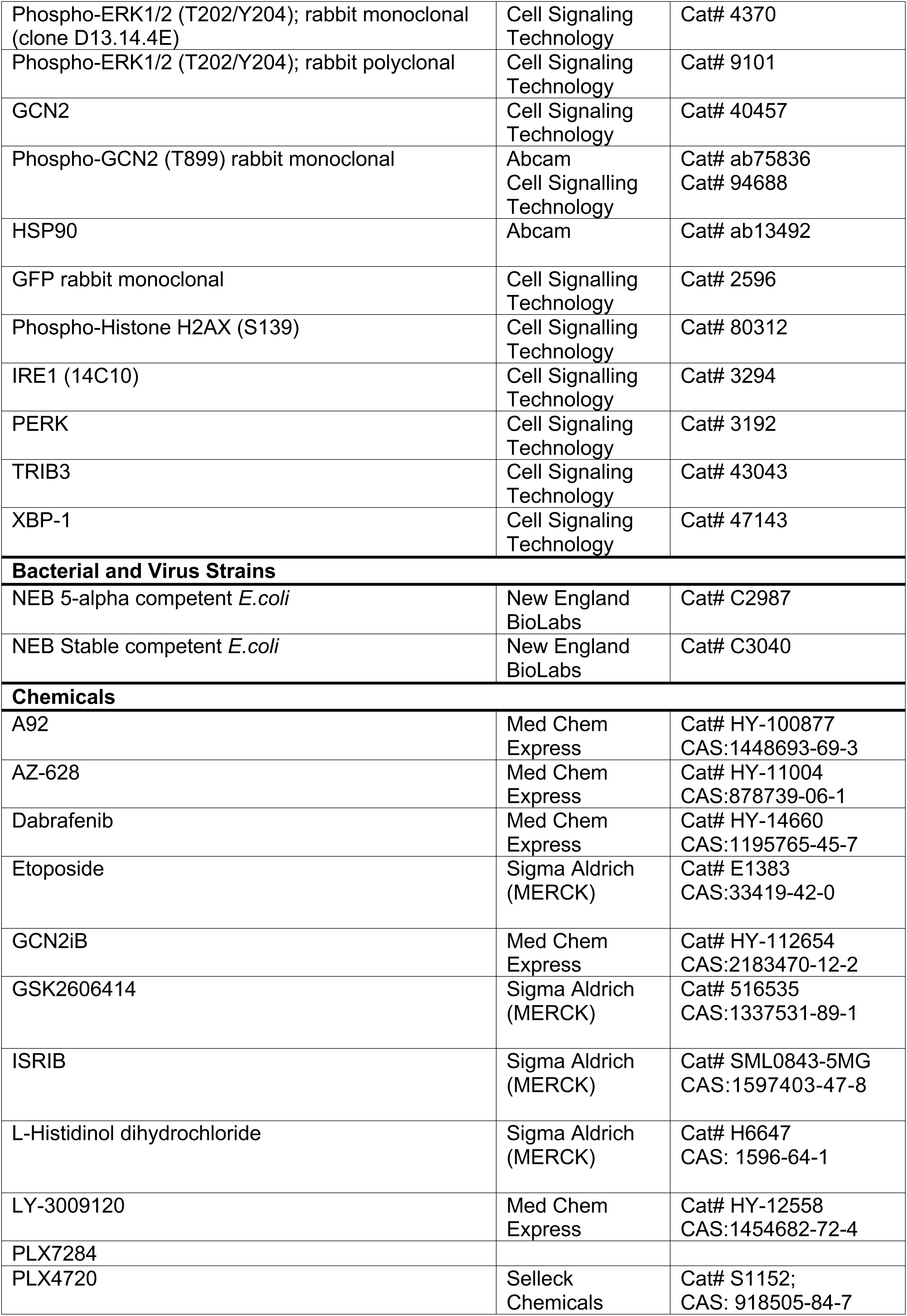
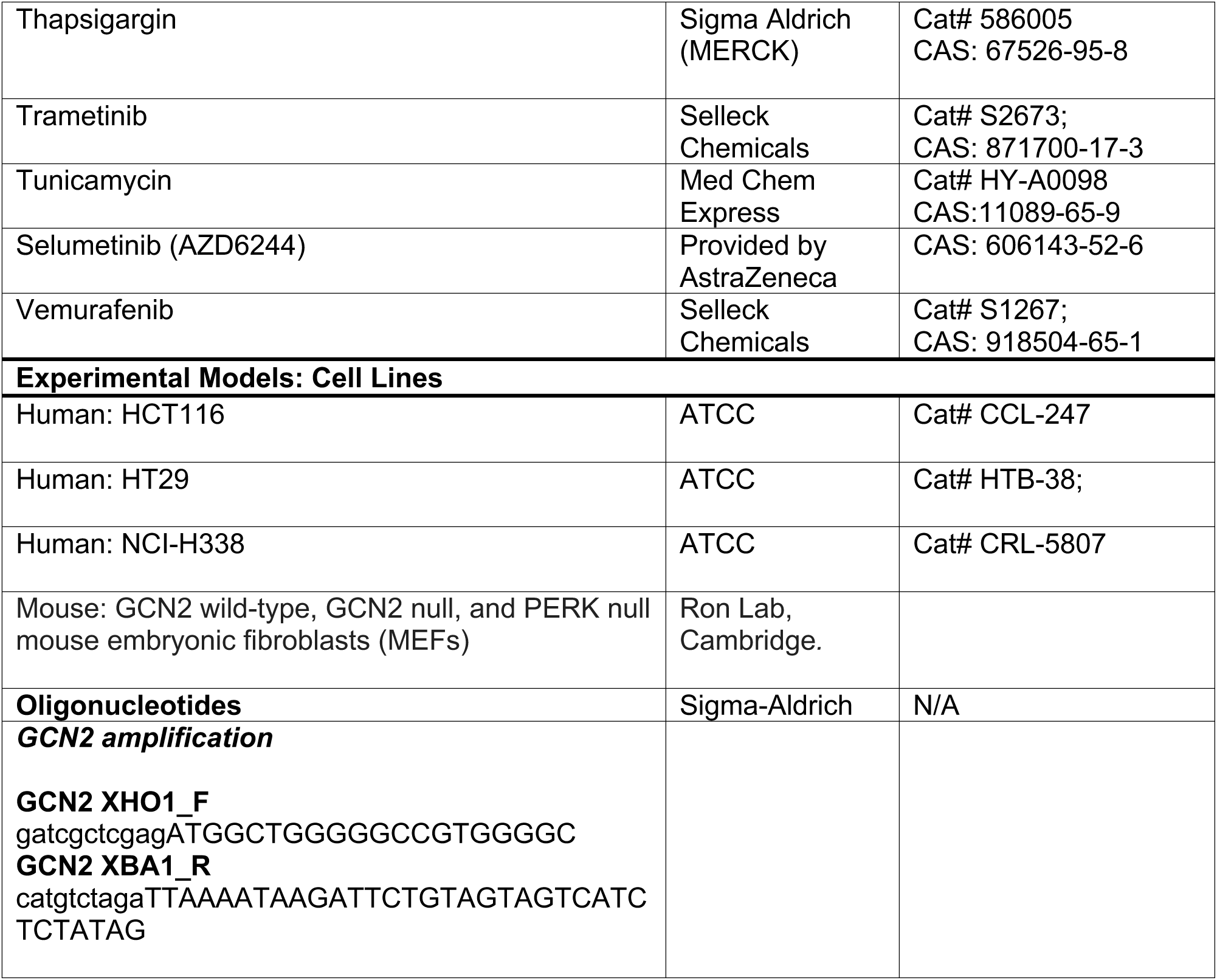
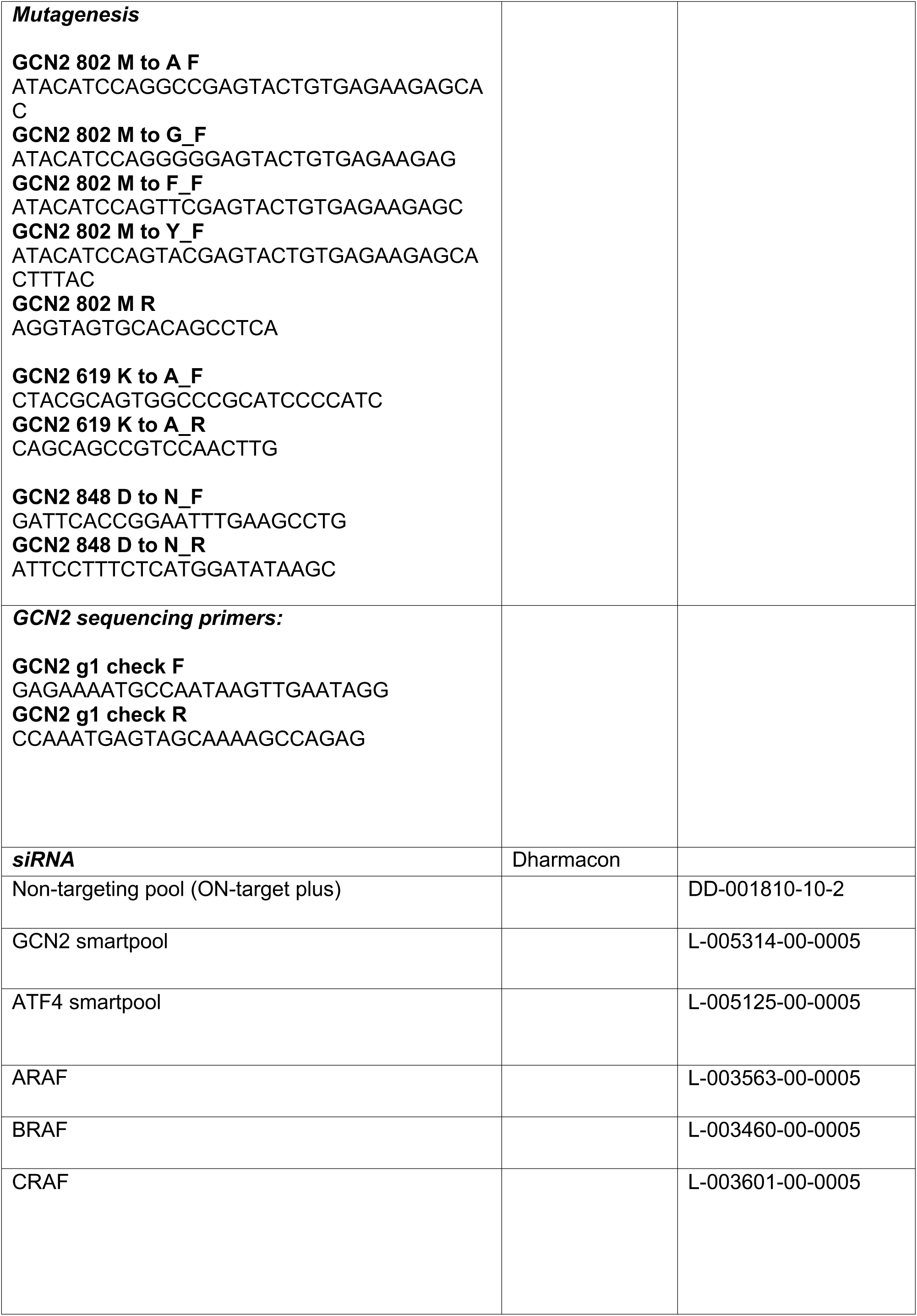
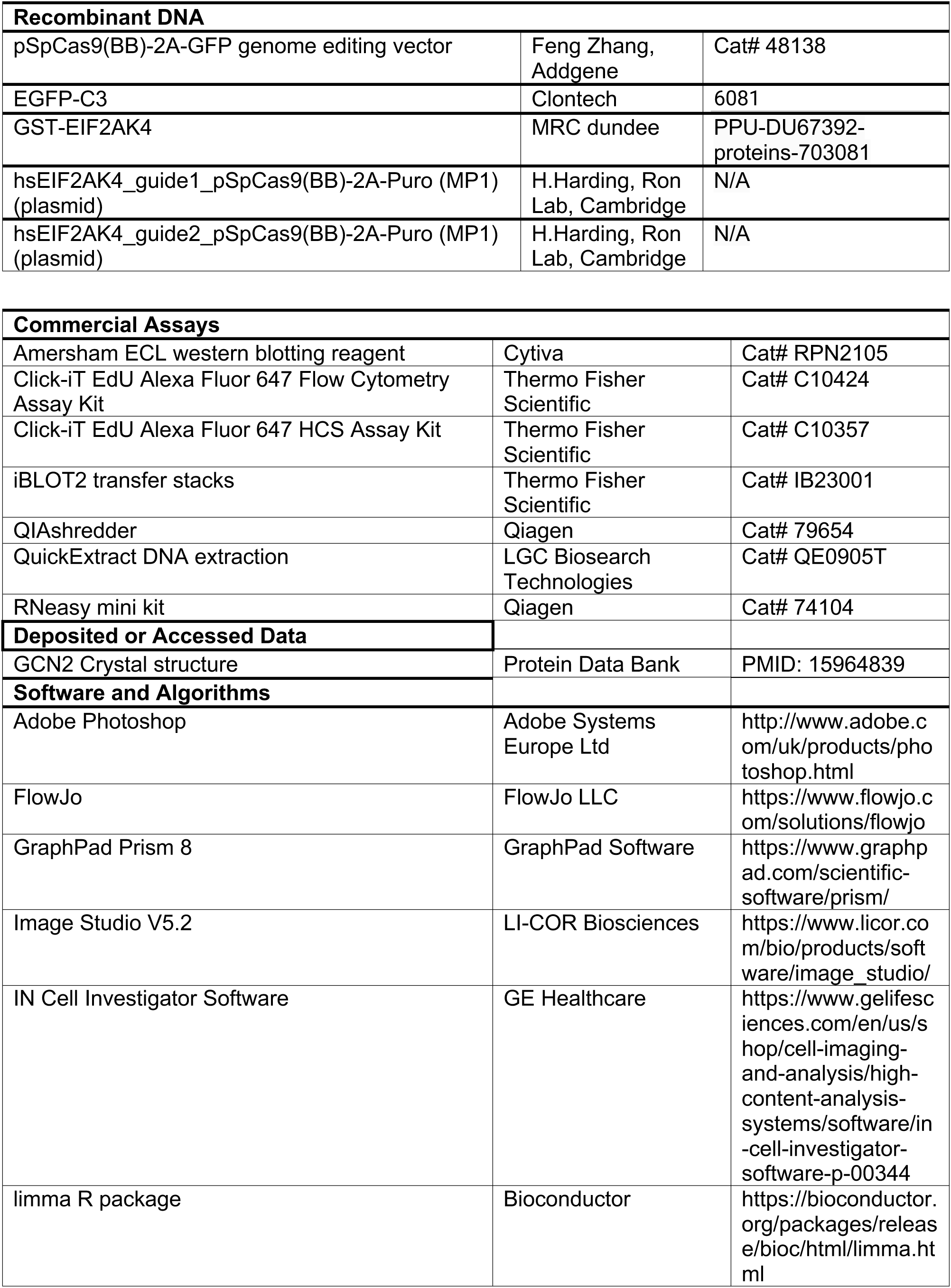
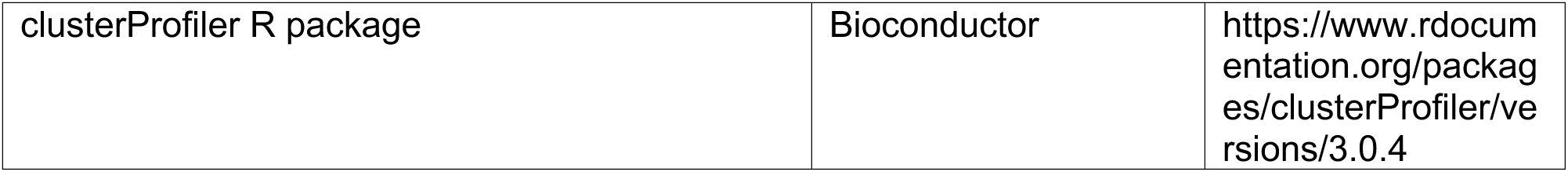
Full list of reagents and resources used in this study.

## References

1. Prior, I.A., Lewis, P.D. and Mattos, C. A comprehensive survey of Ras mutations in cancer. Cancer Res. 72, 2457–2467 (2012)

2. Davies, H., Bignell, G.R., Cox, C., Stephens, P., Edkins, S., Clegg, S. et al. Mutations of the BRAF gene in human cancer. Nature 417, 949–954 (2002)

3. Holderfield, M., Deuker, M.M., McCormick, F. and McMahon, M. Targeting RAF kinases for cancer therapy: BRAF-mutated melanoma and beyond. Nat. Rev. Cancer 14, 455–467 (2014).

4. Caunt, C.J., Sale, M.J., Smith, P.D. and Cook, S.J. MEK1 and MEK2 inhibitors and cancer therapy: the long and winding road. Nat. Rev. Cancer 15, 577–592 (2015).

5. Haling, J.R., Sudhamsu, J., Yen, I., Sideris, S., Sandoval, W., Phung, W. et al. Structure of the BRAF-MEK complex reveals a kinase activity independent role for BRAF in MAPK signaling. Cancer Cell 26, 402–413 (2014).

6. Yao, Z., Torres, N.M., Tao, A., Gao, Y., Luo, L., Li, Q. et al. BRAF mutants evade ERK-dependent feedback by different mechanisms that determine their sensitivity to pharmacologic inhibition. Cancer Cell 28, 370–383 (2015).

7. Bollag, G., Hirth, P., Tsai, J., Zhang, J., Ibrahim, P.N., Cho, H. et al. Clinical efficacy of a RAF inhibitor needs broad target blockade in BRAF-mutant melanoma. Nature 467, 596–599 (2010)

8. Chapman, P.B., Hauschild, A., Robert, C., Haanen, J.B., Ascierto, P., Larkin, J. et al. Improved survival with vemurafenib in melanoma with BRAF V600E mutation. N. Engl. J. Med 364, 2507–2516 (2011).

9. Hauschild, A., Grob, J-J., Demidov, L.V., Jouary, T., Gutzmer, R., Millward, M. et al. Dabrafenib in BRAF-mutated metastatic melanoma: a multicentre, open-label, phase 3 randomised controlled trial. Lancet 380, 358–365 (2012).

10. Robert, C., Karaszewska, B., Schachter, J., Rutkowski, P., Mackiewicz, A., Stroiakovski, D. et al. Improved overall survival in melanoma with combined dabrafenib and trametinib. N. Engl. J. Med. 372, 30–39 (2015).

11. Kopetz, S., Grothey, A., Yaeger, R., Van Cutsem, E., Desai, J., Yoshino, T., et al. Encorafenib, Binimetinib, and Cetuximab in *BRAF* V600E-Mutated Colorectal Cancer N Engl J Med. 381, 1632–1643 (2019).

12. Hall-Jackson CA, Eyers PA, Cohen P, Goedert M, Boyle FT, Hewitt N, et al. Paradoxical activation of Raf by a novel Raf inhibitor. Chem Biol. 6, 559–568 (1999).

13. Durrant, D.E. and Morrison, D.K. Targeting the Raf kinases in human cancer: the Raf dimer dilemma. Brit J. Cancer 118, pages 3–8 (2018)

14. Hatzivassiliou, G., Song, K., Yen, I., Brandhuber, B.J., Anderson, D.J., Alvarado, R. et al. RAF inhibitors prime wild-type RAF to activate the MAPK pathway and enhance growth. Nature 464, 431–435 (2010).

15. Heidorn, S.J., Milagre, C., Whittaker, S., Nourry, A., Niculescu-Duvas, I., Dhomen, N. et al. (2010) Kinase-dead BRAF and oncogenic RAS cooperate to drive tumor progression through CRAF. Cell 140, 209–221 (2010)

16. Poulikakos, P.I., Zhang, C., Bollag, G., Shokat, K.M. and Rosen, N. RAF inhibitors transactivate RAF dimers and ERK signalling in cells with wild-type BRAF. Nature 464, 427–430 (2010).

17. Karoulia, Z., Wu, Y., Ahmed, T.A., Xin, Q., Bollard, J., Krepler, C. et al. An integrated model of RAF inhibitor action predicts inhibitor activity against oncogenic BRAF signaling. Cancer Cell 30, 501–503 (2016).

18. Holderfield M, Nagel TE, Stuart DD. Mechanism and consequences of RAF kinase activation by small-molecule inhibitors. Br J Cancer. 111, 640–645 (2014)

19. Peng, S.B., Henry, J.R., Kaufman, M.D., Lu, W.P., Smith, B.D., Vogeti, S et al. Inhibition of RAF isoforms and active dimers by LY3009120 leads to anti-tumor activities in RAS or BRAF mutant cancers. Cancer Cell 28, 384–398 (2015).

20. Zhang, C., Spevak, W., Zhang, Y., Burton, E.A., Ma, Y., Habets, G. et al. RAF inhibitors that evade paradoxical MAPK pathway activation. Nature 526, 583–586 (2015).

21. Woods, D. Parry D, Cherwinski H, Bosch E, Lees E, McMahon M.et al. Raf-induced proliferation or cell cycle arrest is determined by the level of Raf activity with arrest mediated by p21Cip1. Mol. Cell. Biol. 17, 5598–5611 (1997).

22. Sale, M.J., Balmanno, K., Saxena, J., Ozono, E., Wojdyla, K., McIntyre, R.E. et al. MEK1/2 inhibitor withdrawal reverses acquired resistance driven by BRAF^V600E^ amplification whereas KRAS^G13D^ amplification promotes EMT-chemoresistance. Nat Commun 10(1):2030 (2019)

23. Sale, M.J., Balmanno, K., Cook, S.J. Resistance to ERK1/2 pathway inhibitors; sweet spots, fitness deficits and drug addiction Cancer Drug Resist. 2, 365–380 (2019).

24. Pakos-Zebrucka, K., Koryga, I., Mnich, K., Ljujic, M., Samali, A., Gorman, A.M. The integrated stress response. EMBO Rep. 17, 1374–1395 (2016)

25. Neill, G., Masson, G.R. A stay of execution: ATF4 regulation and potential outcomes for the integrated stress response. Front Mol Neurosci. 16, 1112253 (2023)

26. Kidger, A.M., Munck, J.M., Saini, H.K., Balmanno, K., Minihane, E., Courtin, A. et al. Dual-Mechanism ERK1/2 Inhibitors Exploit a Distinct Binding Mode to Block Phosphorylation and Nuclear Accumulation of ERK1/2 Mol Cancer Ther. 19, 525–539 (2020).

27. Balmanno, K., Kidger, A.M., Byrne, D.P., Sale, M.J., Nassman, N., Eyers, P.A., Cook, S.J. ERK1/2 inhibitors act as monovalent degraders inducing ubiquitylation and proteasome-dependent turnover of ERK2, but not ERK1. Biochem J. 480, 587–605 (2023).

28. Walter, P, and Ron, D. The unfolded protein response: from stress pathway to homeostatic regulation. Science. 334, 1081–1086 (2011)

29. Madden, E., Logue, S.E., Healy, S.J., Manie, S., Samali, A. The role of the unfolded protein response in cancer progression: From oncogenesis to chemoresistance. Biol Cell 111, 1–17 (2019).

30. Eyers, P.A., Keeshan, K., Kannan, N. Tribbles in the 21st Century: The Evolving Roles of Tribbles Pseudokinases in Biology and Disease. Trends Cell Biol. 27, 284–298 (2017).

31. Axten, J.M., Medina, J.R., Feng, Y., Shu, A., Romeril, S.P., Grant, S.W, et al. Discovery of 7-methyl-5-(1-{[3-(trifluoromethyl)phenyl]acetyl}-2,3-dihydro-1H-indol-5-yl)-7H-pyrrolo[2,3-d]pyrimidin-4-amine (GSK2606414), a potent and selective first-in-class inhibitor of protein kinase R (PKR)-like endoplasmic reticulum kinase (PERK). J Med Chem. 55, 7193–207 (2012).

32. Sidrauski, C., McGeachy, A.M., Ingolia, N.T., Walter, P. The small molecule ISRIB reverses the effects of eIF2α phosphorylation on translation and stress granule assembly Elife. 4:e05033 (2015).

33. Zyryanova, A.F., Kashiwagi, K., Rato, C., Harding, H.P., Crespillo-Casado, A., Perera, L.A. et al. ISRIB Blunts the Integrated Stress Response by Allosterically Antagonising the Inhibitory Effect of Phosphorylated eIF2 on eIF2B. Mol Cell. 81, 88–103.e6 (2021).

34. Brazeau, J.F., Rosse, G.. Triazolo[4,5-d]pyrimidine Derivatives as Inhibitors of GCN2. ACS Med Chem Lett. 5, 282–283 (2014)

35. Karaman, M.W., Herrgard, S., Treiber, D.K., Gallant, P., Atteridge, C.E., Campbell, B.T., et al. A quantitative analysis of kinase inhibitor selectivity. Nat Biotechnol. 26,127–32 (2008).

36. Li, Y., Cheng, H., Zhang, Z., Zhuang, X., Luo, J., Long, H., et al. N-(3-Ethynyl-2,4-difluorophenyl)sulfonamide Derivatives as Selective Raf Inhibitors. ACS Med Chem Lett. 6, 543–547 (2015)

37. Fujimoto, J., Kurasawa, O., Takagi, T., Liu, X., Banno, H., Kojima, T., Identification of Novel, Potent, and Orally Available GCN2 Inhibitors with Type I Half Binding Mode. ACS Med Chem Lett. 10, 1498–1503 (2019)

38. Maia de Oliveira, T., Korboukh, V., Caswell, S., Winter Holt, J.J., Lamb, M., Hird, A.W., et al. The structure of human GCN2 reveals a parallel, back-to-back kinase dimer with a plastic DFG activation loop motif. Biochem J. 477, 275–284 (2020).

39. Haling, J.R., Sudhamsu, J., Yen, I., Sideris, S., Sandoval, W., Phung, W. Structure of the BRAF-MEK complex reveals a kinase activity independent role for BRAF in MAPK signaling. Cancer Cell. 26, 402–413 (2014).

40. Narasimhan, J., Staschke, K.A., Wek, R.C. Dimerization is required for activation of eIF2 kinase Gcn2 in response to diverse environmental stress conditions. J Biol Chem. 279, 22820–22832 (2004).

41. Masson, G.R. Towards a model of GCN2 activation. Biochem Soc Trans. 47, 1481–1488 (2019).

42. Padyana, A.K., Qiu, H., Roll-Mecak, A., Hinnebusch, A.G., Burley, S.K. Structural basis for autoinhibition and mutational activation of eukaryotic initiation factor 2alpha protein kinase GCN2. J Biol Chem. 280, 29289–29299 (2005)

43. Bayliss, R., Haq, T., Yeoh, S. The Ys and wherefores of protein kinase autoinhibition. Biochim Biophys Acta. 1854,1586–1594 (2015).

44. Taylor, S.S., Shaw, A..S, Kannan, N., Kornev, A.P. Integration of signaling in the kinome: Architecture and regulation of the αC Helix. Biochim Biophys Acta. 1854, 1567–1574 (2015).

45. Arter, C., Trask, L., Ward, S., Yeoh, S., Bayliss, R. Structural features of the protein kinase domain and targeted binding by small-molecule inhibitors. J Biol Chem. 298(8):102247 (2022).

46. Eyers, P.A., Murphy, J.M. Dawn of the dead: protein pseudokinases signal new adventures in cell biology. Biochem Soc Trans. 41, 969–74 (2013)

47. Bailey, F.P., Andreev, V.I., Eyers, P.A. The resistance tetrad: amino acid hotspots for kinome-wide exploitation of drug-resistant protein kinase alleles. Methods Enzymol. 548,117–146 (2014).

48. Eyers, P.A., Craxton, M., Morrice, N., Cohen, P., Goedert, M. Conversion of SB 203580-insensitive MAP kinase family members to drug-sensitive forms by a single amino-acid substitution. Chem Biol. 5, 321–328 (1998).

49. Whittaker, S., Kirk, R., Hayward, R., Zambon, A., Viros, A., Cantarino, N. et al. Gatekeeper mutations mediate resistance to BRAF-targeted therapies. Sci Transl Med. 2, 35ra41 (2010).

50. Nakamura, A. Nambu, T., Ebara, S. Hasegawa, Y., Toyoshima, K., Tsuchiya, Y., et al. Inhibition of GCN2 sensitizes ASNS-low cancer cells to asparaginase by disrupting the amino acid response. Proc. Natl. Acad. Sci. U. S. A., 115, E7776–E7785 (2018)

51. Li, J.M. and Jin, J. CRL Ubiquitin Ligases and DNA Damage Response. Front Oncol. 2, 29 (2012).

52. Rasmussen, D.M., Semonis, M.M., Greene, J.T., Muretta, J.M., Thompson, A.R., Ramos S.T. et al. Allosteric coupling asymmetry mediates paradoxical activation of BRAF by type II inhibitors. eLife 13,RP95481 (2024).

53. Lavoie, H., Li, J.J., Thevakumaran, N., Therrien, M., Sicheri, F. Dimerization-induced allostery in protein kinase regulation. Trends Biochem Sci. 39, 475–486 (2014).

54. Yeung, W., Ruan, Z., Kannan, N. Emerging roles of the αC-β4 loop in protein kinase structure, function, evolution, and disease. IUBMB Life. 72,1189–1202 (2020).

55. Carlson, K.R., Georgiadis, M.M., Tameire, F., Staschke, K.A., Wek, R.C. Activation of Gcn2 by small molecules designed to be inhibitors. J Biol Chem. 299,104595 (2023).

56. Szaruga, M., Janssen, D.A., de Miguel, C., Hodgson, G., Fatalska, A., Pitera, A.P. et al. Activation of the integrated stress response by inhibitors of its kinases. Nat Commun. 14, 5535 (2023).

57. Neill, G. Vinciauskaite, V., Pau, M., Gilley, R., Cook, S.J. and Masson, G.R. Paradoxical Activation of GCN2 by ATP-competitive inhibitors via structural changes which promotes activating phosphorylation by other GCN2 molecules. In review (2024).

58. Ciudad, M.T., Quevedo, R., Lamorte, S., Jin, R., Nzirorera, N., Koritzinsky, M. Dabrafenib Alters MDSC Differentiation and Function by Activation of GCN2. Cancer Res Commun. 4, 765–784 (2024).

59. Nagasawa, I., Kunimasa, K., Tsukahara, S., Tomida, A. BRAF-mutated cells activate GCN2-mediated integrated stress response as a cytoprotective mechanism in response to vemurafenib. Biochem Biophys Res Commun. 482, 1491–1497 (2017).

60. Tang, C.P., Clark, O., Ferrarone, J.R., Campos, C., Lalani, A.S., Chodera, J.D. GCN2 kinase activation by ATP-competitive kinase inhibitors. Nat Chem Biol. 18, 207–215 (2022).

61. Cook, S.J., Stuart, K., Gilley, R., Sale, M.J. Control of cell death and mitochondrial fission by ERK1/2 MAP kinase signalling. FEBS J. 284, 4177–4195 (2017)

62. Chen, S. Ultrafast one-pass FASTQ data preprocessing, quality control, and deduplication using fastp. iMeta 2, e107 (2023).

63. Patro, R., Duggal, G., Love, M. I., Irizarry, R. A., & Kingsford, C. (2017). Salmon provides fast and bias-aware quantification of transcript expression. Nature Methods. 14, 417–419 (2017)

64. Stolarczyk, M., Reuter, V.P., Smith, J.P, Magee, N.E., Sheffield, N.C. Refgenie: a reference genome resource manager. GigaScience. 9, giz149 (2020).

65. Love, M.I., Soneson, C., Hickey, P.F., Johnson, L.K. Pierce, N.T., Lori Shepherd, L., et al.. Tximeta: Reference sequence checksums for provenance identification in RNA-seq. PLOS Computational Biology. 16, e1007664 (2020).

66. Chen, Y., Chen, L., Lun, A.T.L., Baldoni, P.L., Smyth, G.K. edgeR 4.0: powerful differential analysis of sequencing data with expanded functionality and improved support for small counts and larger datasets. bioRxiv (2024) doi: 10.1101/2024.01.21.576131

67. Ritchie, M.E., Phipson, B., Wu, D., Hu, Y., Law, C.W., Shi, W., and Smyth, G.K. limma powers differential expression analyses for RNA-sequencing and microarray studies. Nucleic Acids Research 43, e47 (2015).

68. Inglis, A.J., Masson, G.R., Shao, S., Perisic, O., McLaughlin, S.H., Hegde, R.S., Williams, R.L. Activation of GCN2 by the ribosomal P-stalk. Proc Natl Acad Sci U S A. 116, 4946–4954 (2019)

69. Kochnev, Y., Hellemann, E., Cassidy, K.C., Durrant, J.D. Webina: an open-source library and web app that runs AutoDock Vina entirely in the web browser. Bioinformatics. 36, 4513–4515 (2020)

70. Modi, V., Dunbrack, R.L. Jr. Kincore: a web resource for structural classification of protein kinases and their inhibitors. Nucleic Acids Res. 50, D654–D664 (2022)

71. Faezov, B., Dunbrack, R.L. Jr. AlphaFold2 models of the active form of all 437 catalytically competent human protein kinase domains. bioRxiv. 2023.07.21.550125. (2023). doi: 10.1101/2023.07.21.550125.

